# Systematic analysis of transcription factor combinatorial binding uncovers TEAD1 as an antagonist of tissue-specific transcription factors in human organogenesis

**DOI:** 10.1101/2023.10.05.561094

**Authors:** Araceli Garcia-Mora, Joshua Mallen, Peyman Zarrineh, Neil Hanley, Dave Gerrard, Nicoletta Bobola

## Abstract

Gene expression is largely controlled by transcription factors (TFs), which bind to enhancers in combination with other TFs in a mechanism known as combinatorial binding. While combinatorial binding is well established, a comprehensive view of tissue-specific TF combinations at active enhancers during human embryonic development is still lacking. Using a two-step pipeline to detect co-occurring TF motifs in developmental enhancers across 11 human embryonic tissues, we found that motifs recognized by ubiquitous TF families, including TEAD, TALE, ETS, and STAT, are enriched near tissue-specific sequence signatures in developmental enhancers across multiple tissues. In human heart enhancers, TEAD and GATA motifs frequently co-occur, and in the developing mouse heart TEAD1 and GATA4 co-occupy a set of genomic regions, which are also preferentially bound by CHD4, a component of the NuRD complex involved in transcriptional repression. Consistently, TEAD1 attenuates enhancer activation in vitro, with this repressive effect dependent on tissue-specific activators. Overall, our findings reveal universal patterns of TF connectivity within organ-specific transcriptional networks and highlight a broad, previously unrecognized role for TEAD in coordinating organ growth and differentiation across multiple tissues.

## INTRODUCTION

Gene expression programs determine and maintain cellular identity and function in embryonic development and adult tissue homeostasis, and their dysregulation causes a broad range of diseases (1). Gene expression is largely controlled by transcription factors (TFs), which bind to distal enhancers (in the linear genome) to facilitate recruitment of RNA Pol II at promoters (2,3).

TFs operate in highly interconnected networks. Examples from animal development or cellular reprogramming indicate that transcriptional regulation is commonly mediated by distinct combinations of TFs (3,4). This combinatorial mechanism allows the integration of multiple biological inputs at cis-regulatory elements, resulting in highly diverse regulatory outputs in space and time (2), as well as precise fine-tuning of gene expression (5). Combinatorial TF binding is closely linked with TF cooperativity, where the binding of one TF increases the likelihood or affinity of another TF binding to a nearby site. Several mechanisms of TF cooperativity have been described. In direct cooperativity TFs interact through direct protein-protein contacts, forming hetero-or homodimers that establish more stable, higher-affinity interactions with DNA (6). In contrast, indirect cooperativity is a less specific mechanism and likely explains the widespread occurrence of combinatorial TF binding on chromatin. In this case, TFs rely on mutual interdependence while engaging with their respective sites on chromatin. Single TFs cannot effectively compete with nucleosomes, but multiple TFs that recognize closely spaced binding sites (within ∼150 bp) synergistically act through ‘mass action’ to displace the same nucleosome. Consequently, they indirectly aid and enhance the binding of each other (6). This extensive cooperativity between TFs at the levels of DNA binding explains why enhancers tend to contain clusters of multiple TF recognition sites.

While the concept of TF combinatorial binding is well established, a systematic exploration of tissue-specific TF combinations during human embryonic development is still missing. Most documented cases of TF combinatorial binding occur in specific contexts, such as distinct cell types or developmental stages. To fully understand how TF synergy shapes transcriptional regulation, it is essential to take a comprehensive view across various tissues and organs. This broader perspective could reveal universal patterns of TF interactions within organ-specific transcriptional networks and enhance our ability to predict the functions of regulatory sequences.

TF combinatorial binding is identified in the genome by the co-occurrence of TF sequence motifs within active regulatory regions. Histones associated with active enhancer regions are often marked by post-translational modifications, notably H3K27ac, which is deposited by p300/CBP enzymes and is predictive of enhancer activity (7). TFs bind DNA in a sequence-specific manner through highly conserved DNA-binding domains. These recognition sequences are represented by position weight matrices (PWMs), which specify the TF’s relative preference for each base at each position in the binding site and enable predictions of TF binding based on DNA sequence. Several methods are available to explore the co-occurrence of TFs at regulatory regions (8–18). Some require ChIP-seq data of target TFs (9,11,16,17) or integrate TF and H3K27ac ChIP-seq experiments (12). These approaches are relatively context-specific as they rely on the occupancy of individual TFs. Other methods do not require TF ChIP-seq experiments, but largely focus on direct cooperativity, promoter regions rather than enhancers or use chromatin accessibility data as input instead of H3K27ac (10,13,14,18).

Here, we systematically investigate TF combinatorial binding at tissue-specific enhancers active during human organogenesis. We designed a two-step pipeline using H3K27ac ChIP-seq and RNA-seq data to identify context-specific, co-occurring TF motifs. This approach reveals both novel and well-established TF interactions. By analyzing the combinatorial TF landscape of developmental enhancers, we found that homotypic motif co-occurrence is a prevalent feature in tissue-specific enhancers. We also identified combinatorial binding of ubiquitous and tissue-specific TFs as a general characteristic of developmental enhancers, with an enrichment of motifs recognized by ubiquitous TF families, including TEAD, TALE, ETS, and STAT, near tissue-specific sequence signatures across multiple tissues. To address the functional role of ubiquitous TFs on organ/cell type-specific transcriptional programs, we focused on the broadly expressed TEAD TFs, identified as combinatorial partner of tissue-specific TFs in heart ventricle, retinal pigmented epithelium (RPE), lower limb, kidney, brain, pancreas and palate. Our results indicate that TEAD1, together with its coactivator YAP, attenuates tissue-specific enhancer activation, pointing at a broad, so far unrecognised effect of TEAD on cell type-specific transcriptional programs. We propose that TEAD may link transcriptional programmes controlling organ growth and differentiation in multiple tissues.

## MATERIAL AND METHODS

### Identification of context-specific co-occurring motifs in human developmental enhancers

#### Human transcriptomic and epigenomic data

RNA-seq data was originally presented in (19) and reanalysed in (20). It is available at www.ebi.ac.uk/ega/home under the accession EGAS00001003738. The RNA-seq data used in this study has been mapped to the hg38 genome. H3K27ac ChIP-seq data was obtained from (20) and is available at www.ebi.ac.uk/ega/home under the accession EGAS0001004335. The genome was parsed into 1kb bins and reads were mapped onto bins. Tissue-specific H3K27ac 1kb regions were identified using a thresholding method based on read counts. Candidate putative tissue-specific enhancers were bins replicated in both replicates of one tissue but not replicated in any other tissue. For details of the method used refer to the original paper (20).

#### Identification of ‘First Search’ TFs expressed during human organogenesis

Motif enrichment analysis of tissue-specific H3K27ac bins (20) was performed for each tissue individually using HOMER’s script findMotifsGenomeWide.pl default settings. The top 20 enriched motifs or motifs enriched with a p-value < 1e^-5^ (whichever came first) were considered.

In parallel, RNA counts across tissues were down-sampled to the 75th percentile. Genes with fewer than 75 reads total across tissues were removed from the analysis and a gene with fewer than 10 reads was considered not expressed in that tissue. Genes encoding TFs (5) were selected and their expression values across tissues was normalised by their median expression. The logarithm of the normalised expression was calculated. To identify tissue-restricted TFs, k-means clustering (k = 10) of TFs was performed using the R package pheatmap (21), and the largest cluster, whose expression remained mostly constant (determined visually) across samples was removed, leaving 9 clusters of TFs with tissue-restricted patterns of expression. For each tissue, TFs with a log normalised expression value > 5 (equivalent to approximately 150 times its median expression) in both replicates were classified as tissue-restricted for that tissue.

#### Clustering PWMs by similarity

The HOMER motif library file (22) was used as a source of PWMs. A clustering method which groups PWMs by distance was used to group all motifs from the HOMER motif library. First, a similarity matrix using the R package universalmotif (23) was calculated for all pairs of PWMs, followed by the calculation of the corresponding distance matrix (1 – similarity matrix). Hierarchical clustering was performed using the base R function hclust, which uses the “complete” agglomeration method as default. A distance tree was plotted, and a cut-off height of 0.5 was used to obtain 79 clusters of motifs. A height of 0.5 was chosen based on the representation of known families. The clusters were named according to TF family systematics (24–28). The most prominent family or subfamily within the group was used. TF subfamilies were used when possible. This was mainly the case for clades of basic Helix-Loop-Helix (bHLH) factors and some Zinc Finger (Zf) subfamilies. In the case where TFs belonging to the same family but spanning across different subfamilies were clustered together, the family and a number were assigned, e.g. Nuclear Receptors (NR) were NR_1, NR_2, NR_3, NR_4, NR_5. Clusters containing only two TFs which belonged to different families were labelled as “other”.

#### Identification of ‘Second Search’ TFs co-occurring with ‘First Search’ TFs

Firstly, the query motif’s (referred to as ‘First Search’ TF motif) PWM was obtained from the HOMER motif library file. Next, the scanMotifGenomeWide.pl script, from HOMER (v4.11, 10-24-2019) (22) was used to obtain a bed file of genomic coordinates belonging to the ‘First Search’ motif in the genome build of interest (e.g. hg38). BEDtools intersect (29) was used to select input and background coordinates for motif enrichment analysis. Input coordinates consist of ‘First Search’ TF motif coordinates that lie within human embryonic H3K27ac-specific 1kb bins. BEDtools random (29) was then used to generate a set of 1kb background regions from the same genome build. The background was filtered to regions containing ‘First Search’ TF motifs. The final background set contained approximately double the number of regions as the foreground set. HOMER uses a binomial distribution to score motifs, which works well when the background sequences outnumber the target sequences, which is why the starting number of random genomic bins is twice the number of tissue-specific bins available (22). Background optimisation was performed before selecting random genomic coordinates as the final background. Backgrounds tested were: 1) random ‘First Search’ TF coordinates as background, 2) non-tissue-specific enhancers consisting of 1kb bins from the human genome acetylated in all tissues studied from (20) 3) HOMER default background consisting of random sequences with matching GC-content distribution as the target sequences and 4) all other putative tissue-specific enhancers from (20). Motif enrichment analysis was performed using the findMotifGenomeWide.pl script, from HOMER using a size parameter of “200”, to search for enrichment motifs 100nt either side of the ‘First Search’ motif coordinates found within context-specific regulatory regions compared to random instances of the “First Search’’ motif. A p-value cut-off of 1×10^-5^ was used and only motifs occurring in 5% or more of foreground regions in HOMER results were considered. Significantly enriched motifs from the HOMER results were substituted by their corresponding cluster of HOMER PWMs. When HOMER results included more than one motif belonging to the same cluster, the motif with lowest p-value was used. The R package ggplot (30) was used to produce bubble plots for network visualization, where negative log p-value determine bubble size. To understand the frequency at which each cluster appeared in the Second Search, a heatmap depicting number of times each cluster appeared as Second Search TF was created using pheatmap (21). Each frequency was normalised by number of motifs in the cluster and the heatmap was scaled by column.

#### Validation of ventricle results with VISTA enhancers

Human experimentally validated heart, limb and forebrain enhancers were downloaded from the VISTA enhancer browser (31). Only enhancers active in exactly one of the three tissues were selected for further analysis resulting in 254 heart enhancers, 400 forebrain enhancers and 258 limb enhancers. Coordinates of ventricle-specific, limb-specific and brain-specific co-occurring motifs predicted using the two-step pipeline described above within each set of VISTA enhancers were identified. This was done by intersecting all instances of the ‘First Search’ motifs in the hg19 genome identified using the HOMER scanMotifGenomeWide.pl script with the VISTA enhancers coordinates using BEDtools intersect. Regions of 100nt either side of the First Search motifs within VISTA enhancers were selected. Finally, these 200nt windows were intersected with all instances of the corresponding ‘Second Search’ TFs, to identify VISTA enhancers containing each pair of predicted ‘First Search’ and ‘Second Search’ TFs within 100nt of each other. The hypergeometric distribution was calculated using the R stats package, to identify whether a pair of co-occurring motifs significantly appeared within 100nt of each other more frequently in one tissue compared to the other two.

#### Validation of GATA4 ventricle results with GATA4 ChIP-seq

An embryonic mouse ventricle GATA4 ChIP-seq (32) (GSE52123) was used to validate predicted functional co-occurring motifs with GATA4 in the human embryonic ventricle (peak calling is described in the following section). Motif enrichment analysis performed on 1) mouse GATA4 200nt peak summits (“peaks”), 2) 100nt either side of GATA4 motifs within mouse GATA4 200nt peak summits (“flanks”), 3) “flanks” within human heart ventricle-specific H3K27ac regions (“H3K27ac”), were compared to the results presented in this manuscript using GATA4 and ventricle-specific H3K27ac bins as input, in other words ventricle ‘Second Search’ results (vSS). For “H3K27ac”, where mouse GATA4 ChIP-seq and human ventricle-specific H3K27ac bins were used, human ChIP-seq peaks were converted to mm10 using the UCSC liftOver conversion tool (33). A heatmap showing the enrichment p-values of motifs enriched within the top 20 known results in “peaks”, “flanks”, “H3K27ac” and “vSS” was created with R base function heatmap. P-values were centred and scaled across rows.

### Functional analysis of TEAD1 using murine data

#### TF ChIP-seq analysis and downstream analysis

Fastq files from TEAD1 mouse adult liver (34) (n=3), TEAD1 mouse embryonic ventricle (35) (n=2), TEAD1 human primary keratinocyte ChIP-seq (36) (n=2), GATA4 mouse E12.5 ventricle ChIP-seq (32) (n=2), GATA6 mouse posterior branchial arch (PBA) ChIP-seq (37) (n=2) and CRX mouse P11 retina ChIP-seq (38) (n=2) were obtained from public databases. Raw fastq files were analysed using an adaptation of the TF ChIP-seq pipeline from (39) for single-end or paired-end. Trimmomatic (40) was used for trimming, Bowtie2 (41) for aligning to the mouse genome (mm10) or human genome (hg38), samtools (42) to remove the aligned reads with a mapping quality of Q10 and MACS2 for peak calling (43) using default narrow peak calling setting for TFs. Blacklisted peaks (44) were removed using BEDtools subtract (29). A cut-off of fold enrichment (FE) > 3 was used to select for peak summits. Windows of 100nt either side of the peak summits were considered as peaks. HOMER (22) was used to check for the enrichment of the corresponding TF motif binding sites to ensure the high quality of the ChIP-seq experiment used. Finally, for the three TEAD1 ChIP-seq experiments, HOMER was used to re-centre the peaks around the corresponding top TEAD1 enriched binding site for each experiment. Any conversions from mm10 to hg38 and vice versa were performed using the UCSC liftOver conversion tool (33). In addition, publicly available bed files of E11.5 mouse ventricle H3K27ac ChIP-seq (n=2) (45) and embryonic mouse ventricle CHD4 ChIP-seq (n=3) (46) were downloaded for analysis. All peak intersections were done using BEDtools intersect (29). R bioconductor packages GenomicRanges (47) and ChIPpeakAnno (48) were used to overlap 200nt peaks around the summits to visualise as Venn diagrams and ChIPseeker (48) was used for peak annotation. For gene ontology (GO) analysis, the Genomic Regions Enrichment of Annotations Tool (GREAT) (49) was used with default parameters. HOMER’s script findMotifsGenomeWide.pl (22) was used for motif enrichment analysis.

#### Identification of candidate enhancers for analysis

To identify candidate putative heart ventricle-specific enhancers bound by GATA6 and TEAD1, mouse PBA GATA6 ChIP-seq summits (37) ventricle GATA4 ChIP-seq summits and (32) ventricle TEAD1 ChIP-seq summits (35) were intersected. Intersecting summits were converted from mm10 to hg38 using the UCSC liftOver conversion tool (33) and intersected with human H3K27ac ventricle-specific 1kb ChIP-seq regions (20) resulting in a list of high-confident ventricle enhancers bound by TEAD1, Gata6 and GATA4. To further reduce the amount of candidate regions, only regions containing GATA4 (50) and TEAD1 (51) motifs (PWM downloaded from HOMER) within 100nt from each other were selected. Finally, the mouse genome informatics database (MGI) (52) was used to identify regions with known cardiovascular system phenotypes. To identify candidate putative RPE-specific enhancers bound by CRX and TEAD1, mouse retina CRX ChIP-seq summits (38) were converted to hg38 and intersected with human H3K27ac RPE-specific 1kb regions (20) To ensure high-confident regions, only regions containing OTX2/CRX (53) motifs were selected. Only regions containing a TEAD1 (38) motif were considered to select final candidate enhancers. Finally, candidate enhancers’ closest genes had either a phenotype or expression profile in the visual system according to the MGI.

### Enhancer cloning

Enhancers were amplified from human genomic DNA using the primers listed in table S1, cloned into pCR8/GW/TOPO vector (Life Technologies) and recombined using the LR Clonase-II Gateway System (ThermoFisher) into pGL4.23-GW (a gift from Jorge Ferrer; Addgene plasmid # 60323; http://n2t.net/addgene:60323RRID:Addgene_60323). Mutant versions of enh_*NKX2.6* and enh_*EYA2* described in table S1 were synthesized by GenScript and cloned as above.

Promoters were amplified from mouse genomic DNA using primers listed in table S1 and cloned into pGL3-Basic (E1751, Promega) using SacI and BglII.

### Cell culture, transfections and luciferase assays

NIH3T3 cells were grown in DMEM (D6429) supplemented with 10% FBS and 5% penicillin/streptomycin and seeded in 24-well plates at 100,000 cells per well. Enhancers were co-transfected with pcDNA3.1 (V790-20, Thermo Fisher), pcDNA3.1(+)-TEAD1 (generated by Genscript), pcDNA-Flag-YAP (18881, Addgene) (54) and pcDNA3-bCrx (bovine Crx, a kind gift from Cheryl M. Craft), pCMV-Gata6 (37) or pcDNA3-GATA4 (55). Cells were transfected with GeneJuice Transfection Reagent (70967, Sigma-Aldrich) 24 hours after seeding, using 250ng of enhancer plasmids and a total of 300ng of expression plasmids per well. Cells were harvested 24 hours after transfection and luciferase measured using Luciferase Assay System and the GloMax Multi-Detection System (Promega). For Verteporfin (VP) experiments, either 0.25uM VP (SML0534-5MG, Sigma-Aldrich) resuspended in DMSO or the equivalent volume of DMSO (Sigma-Aldrich) was added during seeding. More details can be found in table S1.

### Chromatin Immunoprecipitation

Cells were transfected as for luciferase assays, trypsinised, and cross-linked. Cells were lysed by washing with lysis buffer 1 (50mM HEPES, 140mM NaCl, 1mM EDTA, 10% glycerol, 5% NP-40 substitute, 0.25% Triton X-100), followed by lysis buffer 2 (0.2M NaCl, 1mM EDTA, 0.5mM EGTA, 10mM Tris HCl pH 8), and then incubating with lysis buffer 3 (1mM EDTA, 0.5mM EGTA, 10mM Tris HCl pH 8, 0.1M NaCl, 0.1% Na-Deoxycholate, 18.4mM N-lauroyl sarcosine) for 10mins. Nuclei were sonicated in lysis buffer 3 (18 cycles, 30s on and 30s off), Triton X-100 was added to a concentration of 0.1% and the debris was removed by centrifugation. Input sample was collected, and sonication was confirmed by agarose gel. Dynabeads (10002D, Thermo Fisher) were washed (5g/L BSA in PBS pH 7.4) blocked with salmon sperm (15632011, Thermo Fisher) at 0.5mg/ml and bound to antibodies against either CHD4 (ab240640, abcam) or rabbit immunoglobulin G (IgG) (12-370, Sigma-Aldrich), using 5ug per sample (5 million cells). Nuclei lysate was pre-cleared with Dynabeads for 1 hour and then incubated with antibody-bound Dynabeads overnight at 4C° with rotation. Beads were washed (5x 10mins, 4C°, rotating) with cold wash buffer (50mM HEPES, 1mM EDTA, 0.7% Na-Deoxycholate, 1% NP-40 substitute, 0.5M LiCl), cold TE, pH 8 (10mM Tris HCl, 1mM EDTA), and eluted in elution buffer, pH 8 (1% SDS, 50mM Tris HCl, 10mM EDTA) and incubated at 65C°. Eluate was reverse cross-linked at 65C° overnight, treated with proteinase K for 1 hour, and purified using PCR purification kit (28104, QIAGEN). Samples were tested by qPCR. Primers used are listed in table S1. All lysis and wash buffers were kept on ice and contained 1x protease inhibitor.

### Immunofluorescence and microscopy

For IF experiments, NIH3T3 cells were seeded in an eight-chamber slide at 30,000 cells in a final volume of 0.3ml per chamber and fixed after 48 hours. Slides were fixed with 4% PFA for 10 minutes, washed with PBS (3×15mins), and incubated overnight with Purified Mouse Anti-TEF-1 (1:500, BD-610922, BD Biosciences) or mouse anti-YAP1 H-9 (1:150, sc-271134, Santa Cruz) in a PBS solution containing donkey serum albumin and 0.1% Triton. Cells were washed with PBS (3x for 15mins) and treated with Donkey Anti-Mouse IgG H&L (Alexa Fluor 555) (1:500, Ab-150106) and 0.1% Trizol for 2 hours. Cells were stained with 0.5ug/mL DAPI (D9542, Sigma Aldrich) for 5 mins, washed with PBS (2× 5mins), and the slide mounted using Prolong Gold Antifade (ThermoFisher). Experiment was carried out in duplicate. Images were collected on an Axio Imager.M2 Upright microscope (Zeiss) using a 10x objective and captured using a Coolsnap HQ2 camera (Photometrics) through Micromanager software v1.4.23. Specific band pass filter sets for DAPI and Cyf3 were used to prevent bleed through from one channel to the next. Images were analysed using ImageJ (56).

### Statistical analysis

Unless explained otherwise, enhancers were tested in three independent experiments (biological replicates n=3) and three technical replicates per condition (experimental replicates n=3). Verteporfin experiments were tested in two independent experiments (biological replicates n=2) and three technical replicates per condition (experimental replicates n=3). The median of technical replicates was used as a biological replicate, and the average of three biological replicates and the SEM was plotted. Ordinary one-way ANOVA, two-way ANOVA and t-tests were performed using GraphPad Prism version 9.3.1.471 for Windows (www.graphpad.com). Asterisks represent the following values: ns = P > 0.05, * = P < 0.05, ** = P < 0.01, *** = P < 0.001 and **** = P < 0.0001.

## RESULTS

### A pipeline to identify tissue-specific co-occurrence of TF motifs

We designed a two-step pipeline to investigate the combinatorial TF landscape of tissue-specific enhancers. In the first step, we targeted the identification of key events driving acquisition of tissue specificity in distinct tissues. Cell fate and identity hinges on precise gene expression programs directed by lineage-specific TFs. Therefore, our first key step was to identify distinctive sequence signatures linked to individual tissues and their cognate tissue-restricted TFs, or ‘First Search TFs’. This was followed by the identification of TFs potentially cooperating with tissue-restricted TFs, or ‘Second Search TFs’. To identify tissue-specific regulatory regions, we conducted a comparative analysis of H3K27ac ChIP-seq data across 11 embryonic tissues (20). Notably, the H3K27ac histone modification, which marks active enhancers (7), exhibited significantly more distinct and tissue-selective patterns compared to other modifications associated with active transcription. The genome was parsed into 1kb bins and tissue-specific H3K27ac-positive bins were identified using a thresholding method based on read counts. Candidate tissue-specific enhancers were bins present in both replicates of one tissue but not replicated in any other tissue (20). This approach enabled us to isolate and curate developmental enhancers specific to each tissue, whose number varied across the 11 tissues under investigation (Fig 1A). Next, we scanned each set of tissue-specific enhancers for enriched motifs using HOMER (22) and, for each tissue, selected the top 20 signature sequence motifs for further analysis (Fig 1B). To optimise the discovery of tissue-specific regulatory mechanisms, we prioritised cognate TFs based on expression. We analysed tissue-matching transcriptomes to identify tissue-restricted TFs across the developing human embryo. Briefly, RNA-seq counts corresponding to TFs listed in Vaquerizas et al (5) were normalised by median expression across tissues and clustered into tissue-restricted (tissue-specific) and ubiquitous (non-specific) TF clusters using k-means clustering (k = 10) (Fig S1A). The largest cluster, including TFs whose expression levels remained mostly constant (determined visually) across samples was removed, leaving 9 clusters of TFs with tissue-restricted patterns of expression (Fig 1C). Furthermore, to be classified as restricted to a particular tissue, expression of the TF must be approximately 150-fold higher than its median value (Fig 1D, Fig S1B, Table S2). Those TFs demonstrating tissue-restricted expression (as per the method above) and which recognise motifs contained in the top 20 enriched motifs in enhancers active in the same tissue, are catalogued as ‘First Search’ TFs. Due to the similarity in their DNA-binding domains, TFs within the same family often recognize closely resembling motifs. Consequently, our method generated an extensive catalogue of motifs that exhibited significant redundancy. We collapsed all similar motifs in the HOMER motif database and obtained 79 motif clusters by unsupervised classification (Fig S1C, Table S3). If multiple motifs belonging to the same cluster appeared in the top 20 results (Table S3), the PWM motif with the lowest enrichment p-value was used as input. This approach led to the identification of the key transcriptional regulators governing cell fate in individual tissues/organs (Fig 1D, Fig S1B, Table S4). In the example of the heart ventricle, GATA4, HAND1, MYOG (which binds the AP4 motif) and TBX20 were identified as ‘First Search’ TFs (Fig 1D) and their PWM motifs, obtained from the HOMER known motif enrichment analysis results, were used as input for the identification of ‘Second Search’ TF (Fig 1E). In some cases, like in the RPE, multiple ‘First Search’ TFs were represented by the same motif (Table S4).

**Figure 1.**
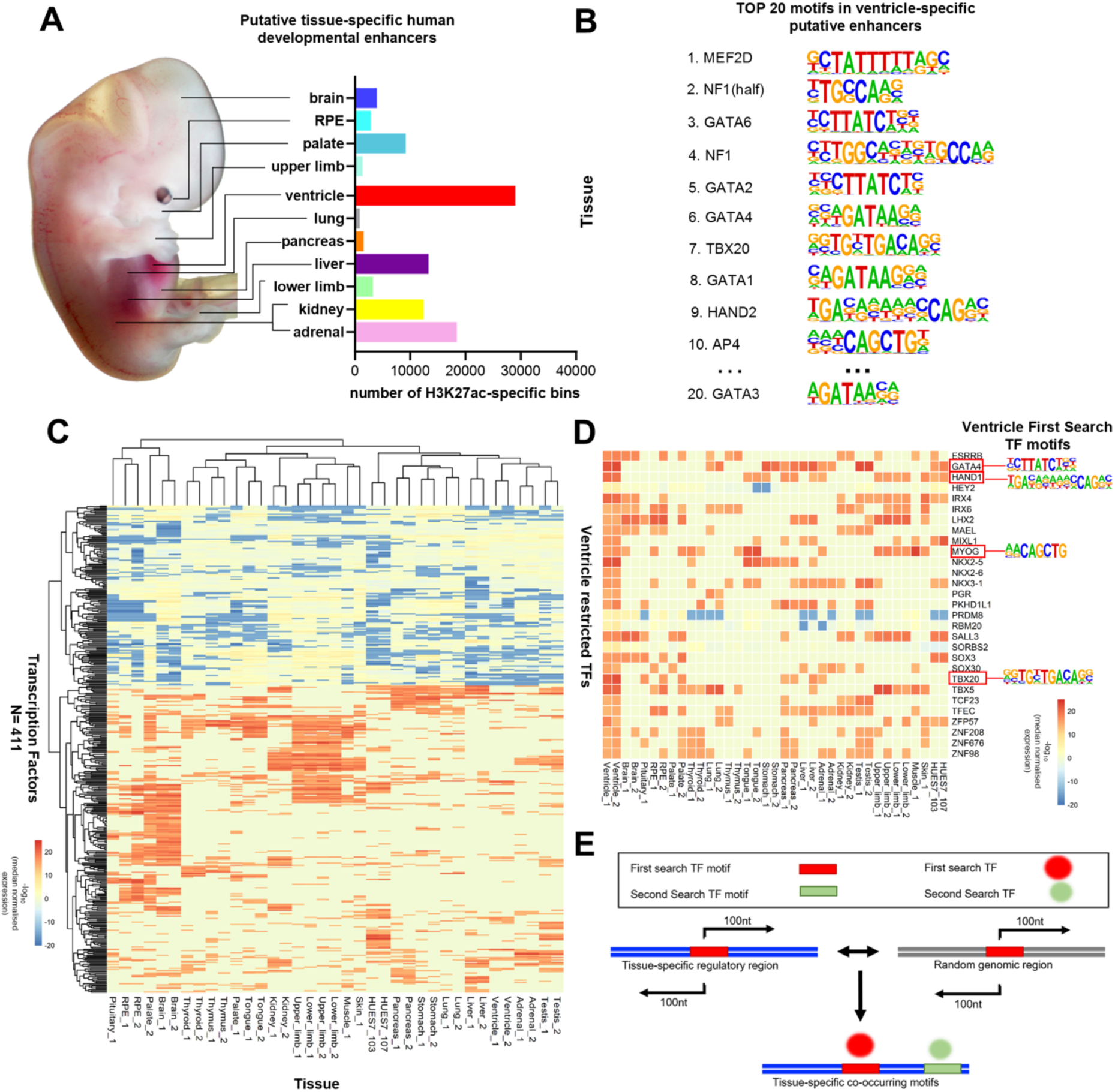
Identification of tissue-restricted TF motifs in human developmental enhancers. A. Human embryo showing tissue/organs used in this study. Samples collected ranged from Carnegie Stage 14 to 21. Black lines connect each organ/tissue to its corresponding number of H3K27ac-positive bins; 1kb bins replicated only in one tissue were used as putative tissue-specific enhancers as described in (20). Each tissue is labelled with the same colour code across the manuscript. B. HOMER known motif enrichment analysis of heart ventricle-specific H3K27ac bins. In this representative example, the top 10 motifs of the list are shown. For analysis, the top 20 known enriched motifs in each set of tissue-specific H3K27ac bins were used. C. Heatmap showing patterns of expression of tissue-restricted TFs across embryonic tissues. All tissues were duplicated except for pituitary, muscle and skin. Raw counts were down sampled based on the 75th percentile and only genes with total expression value over 75 were considered. Expression of each TF was divided by its median expression across all tissues and log median-normalized expression was used to plot the heatmap. TFs whose expression remained constant across tissues were removed. The heatmap of all TFs present in the list obtained from (5) and represented in the RNA-seq data from is shown in figure S1A. D. Heatmap showing the expression of ventricle-restricted TFs. TFs with expression higher than approximately 150 times their median expression in one tissue, were considered restricted to that tissue. Those ventricle-restricted TFs, whose recognition motifs are within the top 20 motif enrichment analysis results shown in 1B were considered ‘First Search TFs’. GATA4, HAND1, MYOG and TBX20 are ventricle-restricted TFs that bind a motif contained in the top 20 enriched motif in ventricle-specific enhancers (1B), namely GATA6, HAND2, AP4 and TBX20 motifs. E. Pipeline to identify ‘Second Search’ TFs co-occurring with ‘First Search’ TFs. Motif enrichment analysis was performed using ‘First Search’ TF motif coordinates within tissue-specific regulatory regions as foreground, and ‘First Search’ TF motif coordinates within random regions as background. Motif enrichment analysis was performed 100nt either side of the ‘First Search’ TF’s motif to identify TF motifs that co-occur with the ‘First Search’ TF motif in tissue-specific regulatory regions. Co-occurring motifs are labelled as ‘Second Search’ TFs.

In a second step, we identified TFs potentially cooperating with tissue-restricted TFs (Fig 1E). For this, we scanned sequences flanking the ‘First Search’ TF motif coordinates in tissue-specific enhancers to identify binding sites for co-binding TFs or ‘Second Search’ TFs. While our ‘First Search’ was tailored to identify tissue-specific regulatory events, the ‘Second Search’ adopted a more permissive approach to accommodate instances of direct and indirect TF binding cooperativity. Specifically, we searched for enriched motifs 100nt either side of the coordinates of the ‘First Search’ motif identified within tissue-specific enhancers, contrasting against random instances of the same motif. We only selected high-confident ‘Second Search’ TF occurrences (p-value cut <1×10^-5^ and foreground motif occurrence > 5%). Significantly enriched motifs derived from the HOMER outcomes were substituted by their corresponding clusters of HOMER PWMs (Fig S1C).

### The network of co-occurring motifs in enhancers active during human organogenesis

Utilizing our two-step pipeline across a diverse spectrum of embryonic tissues provides a panoramic view on the collaborative interactions among TFs. We structured our ‘First’ and ‘Second Search’ outcomes to construct an expansive network delineating the co-occurrence of TF motifs across distinct developmental tissues of the human embryo (Fig 2). The network was built using stringent cut-offs to select highly significant instances of TF motif co-occurrence (foreground enrichment > 5% and p-value < 1e^-5^). We observe well described TF-TF interactions in tissue specification and differentiation in our network, suggesting that our approach is indeed capable of detecting instances of functional TF combinatorial binding. For instance, in the brain, we detected instances of co-binding of basic helix-loop-helix (bHLH) and SOX TFs (ASCL1 and OLIG2 bind a glia-specific enhancer with SOX10 (57)) and of NEUROD and bHLH TFs (NEUROD1 and TCF12 cooperate to drive corticogenesis (58)). Additional examples of functional interactions, present in the network, include GATA4 and HAND2 cooperativity to activate the expression of cardiac-specific genes (59), and TBX20, NKX2.5 and GATA4 interaction during heart development (60). Analysis of liver enhancers identifies the well-established cooperation of FOXA2, GATA4 and HNF4a that primes enhancers for the activation of a liver-specific gene programmes (61,62), as well as co-binding of HNF4A with RXR nuclear receptor (63). Notably, ‘First Search’ TFs belonging to the same tissue do not share the same ‘Second Search’ TFs. This indicates that our two-step approach shows resolution beyond motifs enriched in the entire 1kb bins. For example, in the heart ventricle clusters EBF and RFX_2 only co-occur with TBX20, STAT_1 only co-occurs with GATA4 and ETS only co-occurs with MYOG (Fig 2).

**Figure 2.**
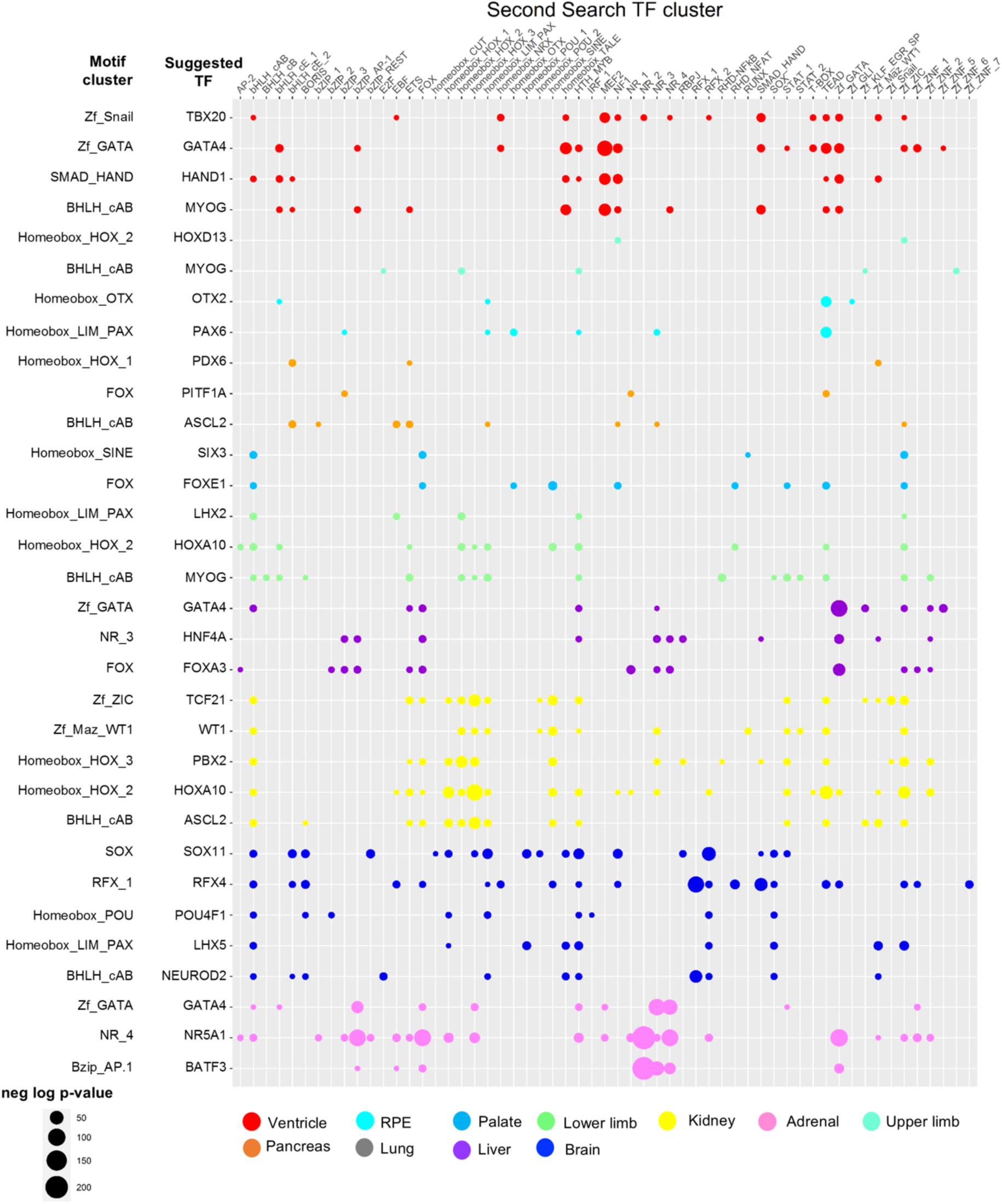
Global network of TF co-binding at tissue-specific developmental enhancers. Network of co-occurring TF motifs at active tissue-specific enhancers in human embryonic development. A cut-off of p-value < 1×10^-5^ and presence in > 5% of foreground regions was used in HOMER known motif enrichment analysis results. On the y-axis are the ‘First Search’ TFs. For each ‘First Search’ TF, its corresponding motif cluster is shown on the left. On the x-axis are TF motif clusters enriched within 100nt either side of the ‘First Search’ TFs (‘Second Search’ TF), clustered by motif similarity (as seen in Fig S1C and table S3). Colours correspond to the ones assigned in Fig 1B. The size of the bubbles matches the negative log p-value: larger bubbles represent smaller p-values.

Our methodology relies on the classification of enhancers based on the presence of H3K27Ac histone modification. Although this histone modification commonly occurs at active enhancer sites, it singularly lacks the ability to conclusively denote functional enhancers, i.e. enhancers that activate gene expression. Consequently, our conclusions might be influenced by the potential misclassification of H3K27Ac-positive sites as functional enhancers when employing H3K27Ac markers alone. To determine the reliability of our results, we examined the occurrence of ‘First’ and ‘Second Search’ TFs within experimentally validated enhancers sourced from the VISTA database (31). We selected heart, forebrain and limb enhancers and performed hypergeometric tests to assess whether ventricle-specific motif pairs identified in our network co-occurred within 100nt of each other significantly more often in heart enhancers, compared to forebrain and limb enhancers. We focused on the following predictions identified in our network: GATA-TEAD, GATA-MEF2, GATA-TBX (TBX clustered within the Zf Snail group) and GATA-NKX. We also used the brain prediction bHLH-RFX, and the limb prediction HOX-PITXEBOX (Pitx1:Ebox clustered within the BHLH_cAB cluster). As expected, GATA-TEAD, GATA-MEF2 and GATA-NKX pairs (co-occurrence within 100nt) were significantly enriched in heart enhancers, compared to limb and brain enhancers (p-value < 0.05) (Fig S2A). In addition, we found the limb signature pair HOX-PITXEBOX, significantly enriched in the limb compared to brain and heart enhancers. We could not detect a significant enrichment of bHLH-RFX in brain enhancers compared to the heart and limb enhancers (Fig S2A). This could be explained by the observation that the bHLH motif CAGCTG is enriched in active enhancers across most tissues (Fig 2).

As an additional validation of our findings, we searched for ‘Second Search’ TFs using experimentally determined TF binding sites. We compared motif enrichment analysis of GATA4 ChIP-seq peaks in mouse embryonic heart (32) with heart ventricle ‘Second Search’ TF results (vSS), using GATA as ‘First Search’ TF. We generated three datasets from GATA4 ChIP-seq: GATA4 ‘peaks’ (200nt centered on the peak summit); GATA4 ‘flanks’ (200nt sequences flanking any GATA4 motifs contained in GATA4 ‘peaks’); and ‘H3K27ac’ (GATA4 ‘flanks’ that also overlap the set of human ventricle-specific enhancers converted to mouse coordinates). Using motif enrichment analysis, we observed a significant overlap between the top ten enriched motifs present in ‘peaks,’ ‘flanks,’ and ‘H3K27ac’, and ‘vSS’ (Fig S2B). The heatmap shows negative log p-values for the top significantly enriched motifs in the four datasets (TEAD, MEF2, TALE, TBX, GATA). TEAD is the top co-occurring motif in ‘peaks’ and ‘flanks’, while MEF2 is the top motif in ‘vSS’ and ‘H3K27ac’. As MEF2 also appears as the top motif in ‘H3K27ac’, this discrepancy is potentially explained by a higher enrichment of GATA-MEF2 sites in active enhancers (relative to all GATA4 binding events). In sum, the analysis of GATA4 occupancy in the heart confirms that GATA4 preferentially bind with MEF2, TALE, TBX, and TEAD, with a tendency for GATA4 and MEF2 to co-occupy active ventricle-specific enhancers. These observations, combined with the identification of known cases of TF cooperativity, suggest that our network provides highly reliable predictions of functional combinatorial binding at tissue-specific enhancers.

Amongst the predictions of our two-step pipeline, we find that sequences surrounding GATA and Forkhead (FOX) motifs in liver enhancers (assigned to GATA4/GATA6 and FOXA3/FOXA2 respectively) are enriched in largely distinct motif clusters. FOX motifs, but not GATA motifs, co-occur with basic leucine-zipper (bZIP) motifs (recognized by ATF, JUN, CEBP and DMRT), and Nuclear Receptor (NR) motifs (particularly clusters NR_1 and NR_4) (Fig S2C). We observe GATA recognition motif, but not FOX motif, to co-occur with zinc finger (ZF) motif clusters, such as those recognized by GLI, KLF, SP, MAZ and ZIC and with BHLH Ebox motifs (BHLH_cAB) (Fig S2C). The observation that GATA4 and FOXA2 motifs in liver enhancers are flanked by distinct collections of sequence motifs suggests that GATA4 and FOXA bind to different groups of liver enhancers. Corroborating this hypothesis, the expression patterns of *Gata4* and *Foxa2* transcripts exhibit only a partial overlap in mouse liver development: *Gata4* expression precedes liver specification, whereas *Foxa2* expression persists throughout hepatic differentiation (64). There is contrasting evidence about the role of the AP-1 complex (formed by TFs such as JUN) as a pioneer TF in the liver (65,66). We find AP-1 motifs to significantly co-occur with FOXA motifs, supporting the hypothesis that AP-1 does not have inherent pioneer abilities (65) and that its apparent pioneer abilities could be due to co-binding with FOXA, a well-established family of liver pioneer factors (67).

### Regulatory patterns in human tissue-specific developmental enhancers

Our panoramic overview of TF combinatorial binding in diverse embryonic tissues offers a unique opportunity to identify universal trends and common patterns of TF interactions within developmental enhancers. By analysing the combinatorial TF landscape of developmental enhancers, we identified homotypic motif co-occurrence as a prevalent feature in tissue-specific developmental enhancers. Homotypic motif pairs predominantly occur across diverse TF families, namely bHLH, AP-1, SOX, ZIC, FOX, HOX, GATA, PAX, NR and Regulatory Factor X (RFX) families (Fig S2D). While TFs belonging to most of these families can bind DNA as homo-or hetero-dimers (68–73), these findings are also consistent with previous studies on enhancer grammar, which have reported homotypic co-occurring motifs to be common, and to occur independently of dimerization events (74,75).

We explored if organ-specific transcriptional networks share patterns of TF functional connectivity. One way to address this is to quantify the occurrence of ‘Second Search’ TF clusters among tissue-specific developmental enhancers. A heatmap of counts of all the ‘Second Search’ TF sites identified in human tissue-specific enhancers (Fig 3A) clustered ‘Second Search’’ TF motifs in three groups: high co-occurring, medium co-occurring and low co-occurring. We found that TFs belonging to the bHLH family are the most frequent ‘Second Search’ TFs in developmental enhancers (Fig 3A). E-box motifs (consensus sequence CAGCTG belonging to the cluster bHLH_C_A_B) are found within 100 nt of the ‘First Search’ TFs in developmental enhancers in eight out of the eleven tissues studied (Fig 2). BHLH TFs play critical roles in the establishment of tissue– and cell-identity in a wide range of tissues (76–81). While these observations highlight a widespread cooperativity of bHLH TFs with tissue-restricted TFs as a key feature of human developmental enhancers, the large size of the bHLH family (> 100 members) can also contribute to the observed high connectivity of BHLH TFs.

**Figure 3.**
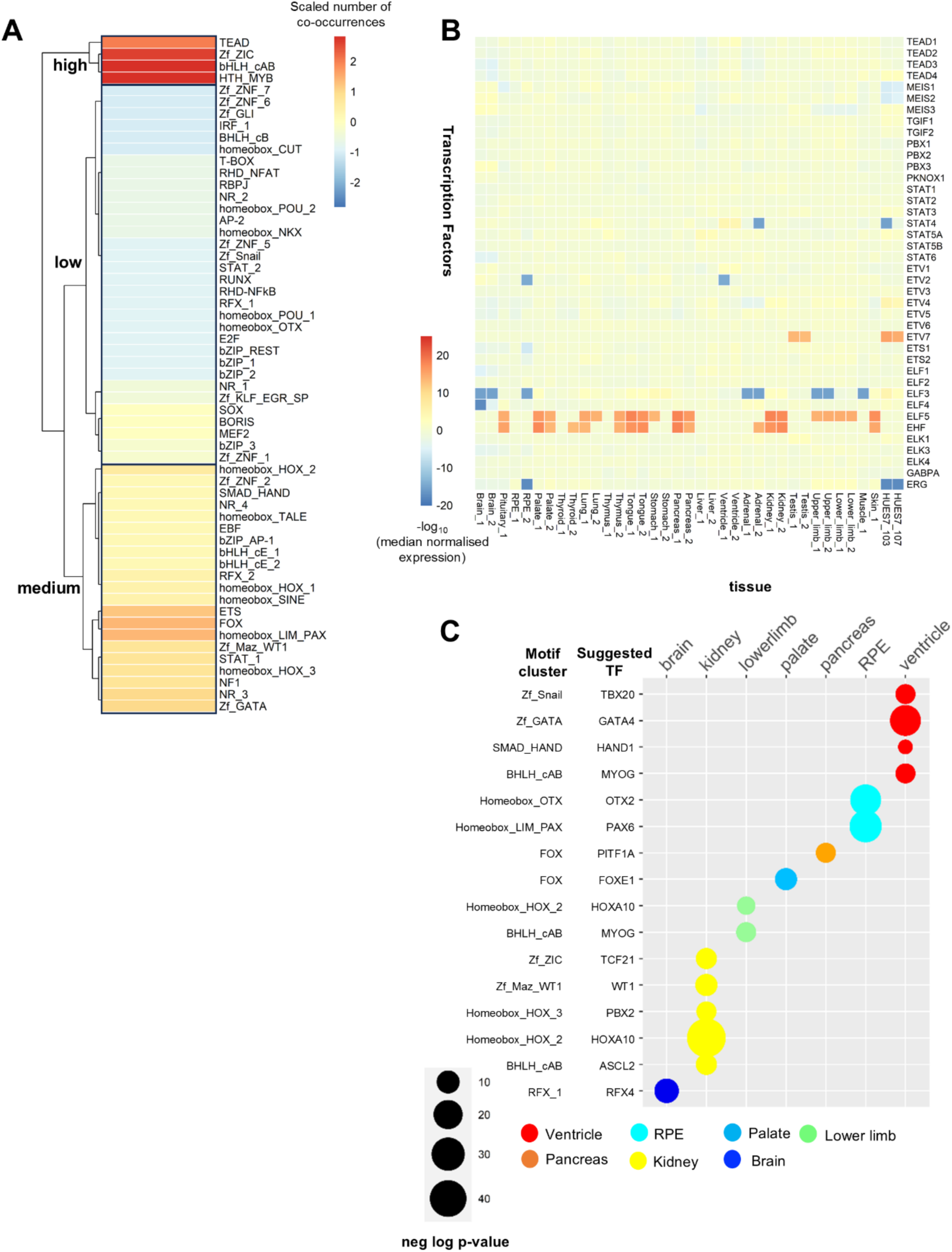
Motifs belonging to ubiquitous families of TFs co-occur with tissue-restricted TF motifs. A. Heatmap showing number of co-occurring events for each ‘Second Search’ TF cluster. Clustering by number of co-occurrences assigns ‘Second Search’ clusters to three groups: high co-occurring, medium co-occurring and low co-occurring. bHLH clades A and B have the highest number of co-occurring events. ‘Second Search’ clusters of motifs recognised by ubiquitous TFs are abundant (found in the high/medium co-occurring groups). For instance, TEAD cluster presents high levels of co-occurring events, whilst TALE and ETS are found in the medium co-occurring cluster. B. Heatmap showing patterns of expression of TFs belonging to TEAD, STAT, ETS and TALE TF families. All tissues were duplicated except for pituitary, muscle and skin. Raw counts were down sampled based on the 75th percentile. Expression of each TF was divided by its median expression across all tissues and log median-normalized expression was used to plot the heatmap. Expression of ubiquitous TFs belonging to the TEAD, TALE, STAT and ETS families do not deviate from their median expression across tissues (with the exception of STAT4, ETV2, ELF3, ERG, ELF5 and EHF). C. ‘First Search’ TFs (y-axis) co-occurring with TEAD binding sites in the tissues shown (x-axis). Colours correspond to the ones assigned in Fig 1B. The size of the bubbles matches the negative log p-value: larger bubbles represented smaller p-values.

Tissue– and cell type-specific patterns of expression can be indicators of TF functions (5,82). TFs can be classified into two groups based on their expression patterns: ubiquitous TFs and tissue-restricted TFs (5). For our analysis, we used hierarchical clustering (k = 10) to distinguish between 9 clusters of TFs with variable expression across tissues and one larger cluster containing ubiquitous TFs whose expression remain constant across tissues (Fig S1A). We identify enrichment of motifs recognised by ubiquitous TF families, TEAD, TALE, ETS and STAT, proximal to tissue-specific sequence signatures across multiple tissues (Fig 3AB). Notably, the TEAD family is classified in the high co-occurrence group, while the other three are in the medium co-occurrence group (Fig 3A). TEAD recognition motifs are significantly enriched in the vicinity of tissue-specific sequence signatures in ventricle, lower limbs, brain, kidney, RPE, pancreas and palate (Fig 3C). TEAD binding sites occur in combination with sixteen ‘First Search’ TFs (Fig 3C). Five of the ‘First Search’ TFs recognise a homeobox cluster motif, suggesting that TEAD TFs widely co-bind with homeodomain TFs. In sum, motifs recognized by families of broadly expressed TFs form a central component of the landscape of tissue-specific developmental enhancers.

### TEAD1 binds tissue– and cell type-specific enhancers

We focused on TEAD TFs to explore the functional relevance of ubiquitous TF binding at tissue-specific enhancers. The small size of the TEAD family, comprising only four members, TEAD1-4 (83), simplifies PWM assignments. In addition, the high number of potential interactions involving TEAD suggests a broad and significant role of TEAD at developmental enhancers. As a first step, we investigated if ubiquitously expressed TEAD TFs bind tissue-specific enhancers. We compared TEAD1 binding in adult mouse liver (34), mouse embryonic ventricle (35) and human primary keratinocytes (36). Overlap of high-confident TEAD1 peaks, centred on their top TEAD motif (Fig 4A), shows that most TEAD1 binding is tissue-or cell type-specific (Fig 4A). While the majority of tissue-specific TEAD1 peaks occur at intronic or distal intergenic regions, as expected for enhancer regions (Fig 4B), a high proportion of overlapping (non-tissue-specific; n=440) TEAD1 peaks occur at promoters. Tissue-specific TEAD1 peaks are associated with tissue-specific biological processes and mouse phenotypes (Fig 4C-E). In contrast, shared TEAD peaks are associated with the hippo signalling pathway and with tumorigenic mouse phenotypes (Fig 4F), consistent with the well-established role of TEAD, and its coactivator YAP, in the hippo pathway and in cancer progression (84,85). Moreover, tissue-specific peaks are enriched in motifs recognised by tissue-specific TFs, such as NKX and TBOX in ventricle peaks (Fig S3A-C), matching our predictions of TEAD binding partners. Compared to tissue-specific peaks, shared non-tissue-specific TEAD1 peaks are enriched in high affinity TEAD site, suggesting that shared peaks may contain higher affinity versions of TEAD binding sites or multiple TEAD motifs (Fig S3D). Overall, these results support our network predictions and suggest that TEAD act in concert with tissue-specific TFs at tissue-specific enhancers.

**Figure 4.**
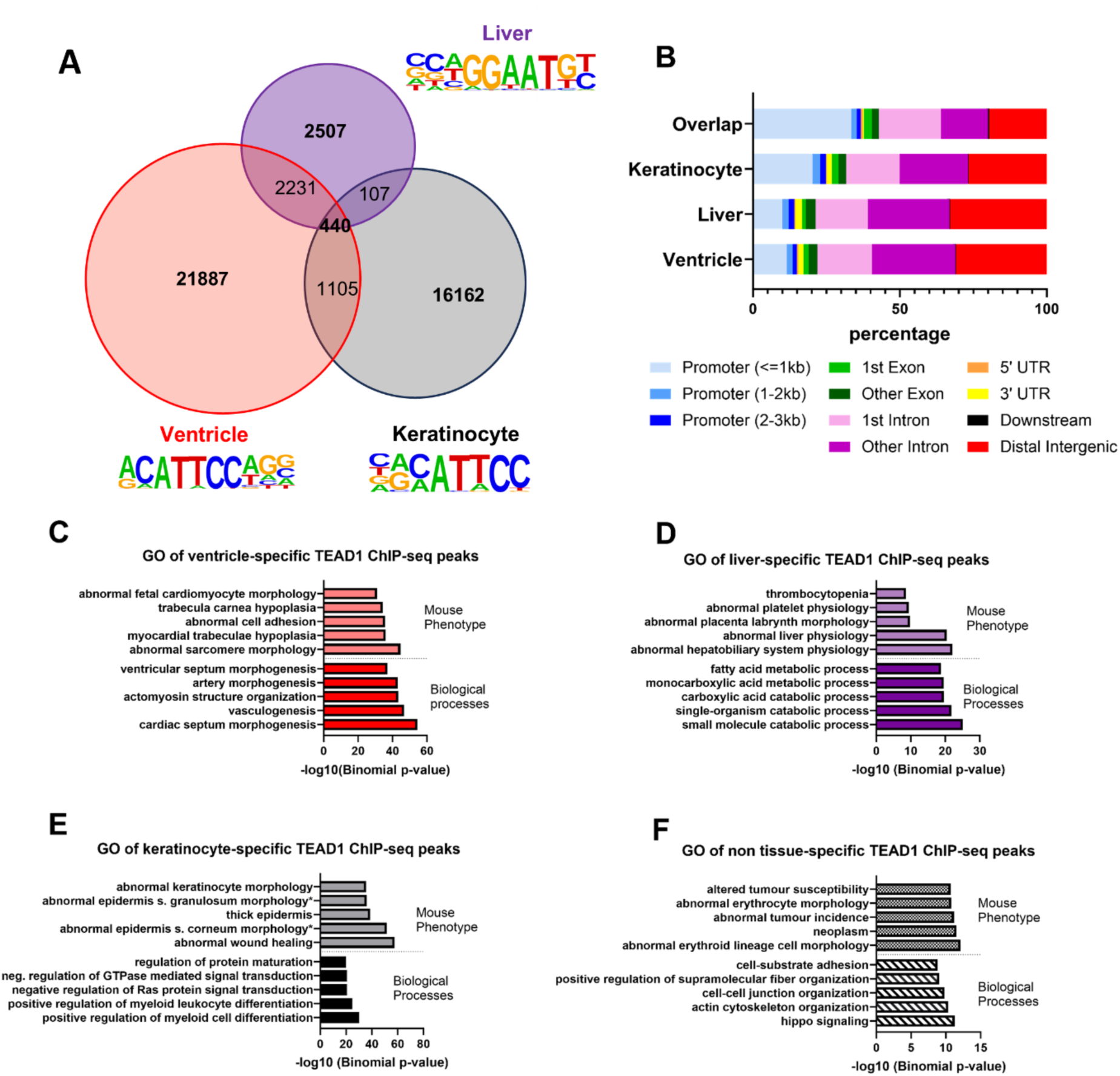
TEAD1 binding occurs mainly at tissue-specific intronic or distal intergenic regions. A. Intersection of adult mouse liver, embryonic mouse ventricle and human primary keratinocyte TEAD1 ChIP-seq shows a high percentage of tissue-specific binding. TEAD HOMER de novo motifs enriched in liver, ventricle and keratinocyte are shown. Each motif was used to centre the corresponding ChIP-seq 200nt regions around the peak summits (referred to as ‘TEAD1 peak’ in the result section). B. Annotation of TEAD1 specific peaks in mouse liver, ventricle and primary keratinocytes and overlapping TEAD1 peaks in all three ChIP-seq experiments. Non-tissue-specific peaks contain a higher percentage of promoters compared to tissue-or cell-type specific TEAD1 binding events. C-F. Top five mouse phenotype and biological process GOs associated with ventricle-specific (C), liver-specific (D), keratinocyte-specific (E) and non-tissue-specific (F) TEAD1 peaks, using the whole mouse genome as background.

### TEAD1 attenuates tissue-specific enhancer activation

While TEAD1 motifs are enriched in active developmental human enhancers, motif discovery alone cannot confirm whether TEAD binds to these enhancers during their active state. To address this, we overlapped TEAD1 binding (E12.5 mouse heart ventricle) to chromatin acetylation (E11.5 mouse heart H3K27ac ChIP-seq) (45) (Fig 5A). We discovered that around 70% of TEAD1 binding is not acetylated (Fig 5A). To investigate the functional significance of TEAD co-binding with tissue-restricted TFs, we performed de novo motif searches on both acetylated and non-acetylated TEAD1 peaks. Our results show that motifs recognized by cardiac TFs co-binding with TEAD in our network (such as TBX, HAND, GATA, MYOG) are more enriched in non-acetylated TEAD1 peaks (Fig 5B and S4AB). Conversely, acetylated TEAD regions are more enriched in motifs recognised by ubiquitous TFs like TALE TF MEIS and MEF2, which are not associated to TEAD motifs in our human developmental enhancers (Fig 5B and S4B). These findings suggest that TEAD1 may co-bind with cardiac-restricted TFs at non-active enhancers, which appears to contradict our identification of TEAD combinatorial binding in acetylated human enhancers. However, it is important to consider that our ventricle-specific enhancers were sampled at a single time point and likely represent enhancers that are selectively active in different cell types and developmental stages, given the heterogeneity of cell types in the ventricle. NKX2-5, TBX5 and GATA4, whose motifs are enriched in TEAD1 non-acetylated peaks, recruit CHD4, a core component of the NuRD complex, to developmental enhancers to suppress non-cardiac cell fates (46). Notably, YAP, a well-established coactivator of TEAD1 (83), directly recruits CHD4 in different contexts, and this recruitment requires TEAD (86). Together these observations suggest that TEAD, in concert with cardiac-restricted transcriptional regulators, may contribute to the recruitment of CHD4. Supporting this hypothesis, the subset of NKX2-5, TBX5 and GATA4 peaks that overlaps with CHD4 binding in the mouse embryonic heart shows a strong enrichment for TEAD motifs (46). In contrast, the fraction of NKX2-5, TBX5 and GATA4 peaks that do not overlap with CHD4 lacks this enrichment (46). Indeed, when overlapping TEAD1 (35) and CHD4 (87) occupancy in the mouse embryonic heart, we found that shared TEAD1-CHD4 peaks are enriched in GATA motifs, while peaks exclusive to TEAD1 are not (Fig 5C). In line with this observation, the intersection of GATA4, TEAD1 and CHD4 occupancy in the mouse heart (Fig 5D) shows that GATA4 occupies the majority (69%) of peaks bound by TEAD1 with CHD4 compared to TEAD1-only peaks (32% overlap) (Fig 5E). Regions co-occupied by GATA4-TEAD1 and CHD4 are associated with cardiomyocyte differentiation (such as cardiocyte differentiation and cardiac muscle cell differentiation) (Fig 5F). Conversely, regions occupied by GATA4–TEAD1 without CHD4 are associated with vasculogenesis (Fig 5G). Collectively, the preferential co-occupancy of GATA4 and TEAD1 at non-acetylated regions bound by CHD4, suggests that TEAD, alongside GATA, may play a role in attenuating the activity of enhancers involved in cardiomyocyte differentiation.

**Figure 5.**
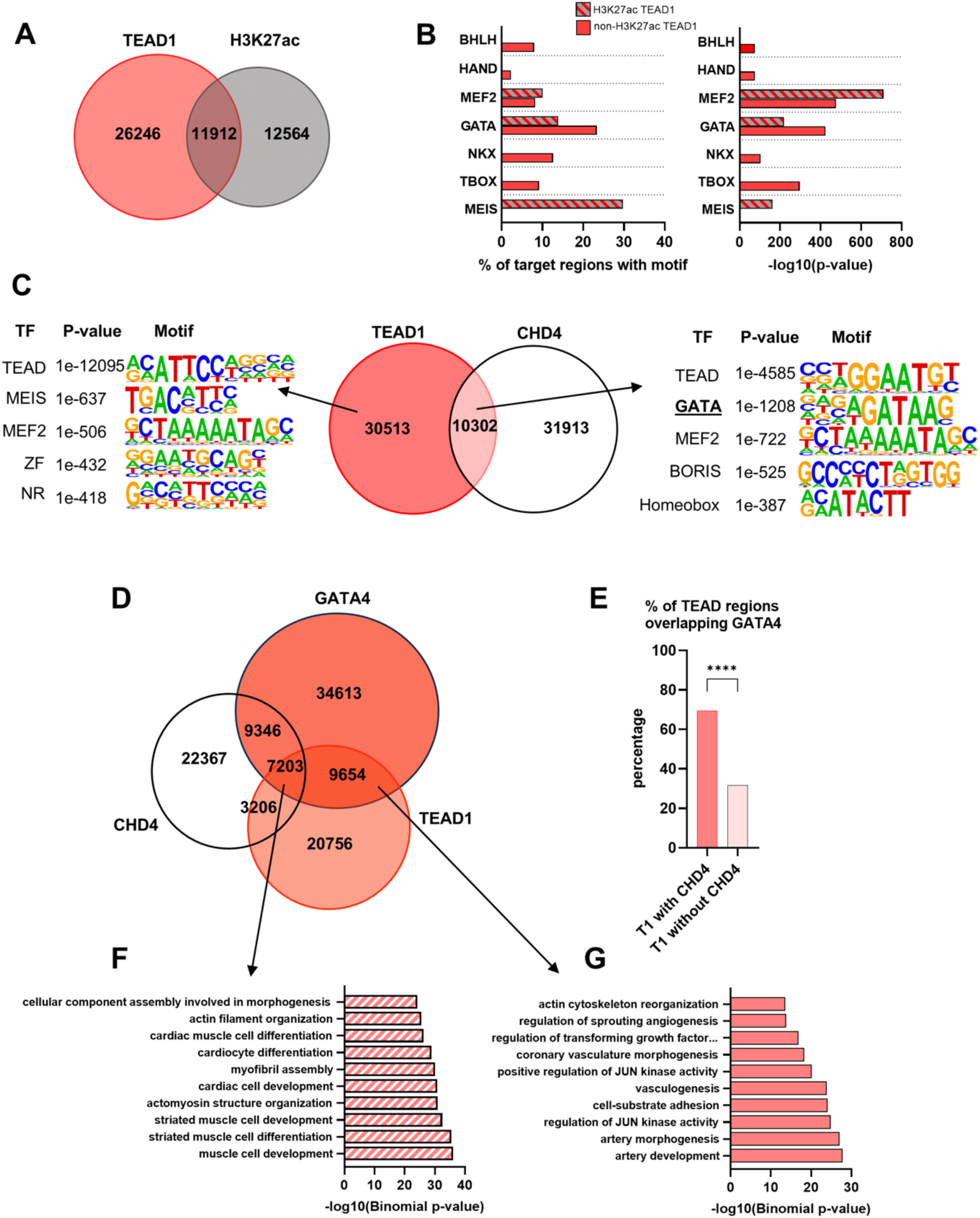
TEAD1 binds regions co-occupied by GATA4 with CHD4 during heart development. A. Overlap between TEAD1 and H3K27ac ChIP-seq peaks in the mouse embryonic heart. B. Sequence motif enrichment in TEAD1 peaks overlapping with H3K27ac (red and grey) and without H3K27ac (red only). The left bar chart shows the percentage of the target regions with motif; the right bar chart shows the neg log p-values of the motif, as identified by HOMER de novo motif enrichment analysis. Cardiac ‘First Search’ TFs and other known cardiac TFs are shown. Motifs bound by tissue restricted TFs HAND, BHLH (Ebox), TBOX and GATA are more enriched in non-H3K27ac TEAD peaks, whilst TF motifs MEF2 and MEIS are preferentially enriched in TEAD1-H3K27ac regions. C. Overlap between TEAD1 and CHD4 ChIP-seq peaks in the embryonic heart ventricle. Top 5 motifs, identified by HOMER de novo motif analysis, are shown for TEAD1 peaks with and without CHD4. GATA motifs are exclusively enriched in TEAD1 peaks overlapping CHD4 peaks. D. Overlap between GATA4, TEAD1 and CHD4 ChIP-seq peaks in the mouse embryonic heart. E. Percentage of TEAD1 peaks overlapping GATA4 peaks. GATA4 preferentially occupies TEAD1-CHD4 regions compared to TEAD1-only regions. Statistical significance was calculated using a two-proportion z test. F. Top ten biological process GOs associated with TEAD1, GATA4 and CHD4 co-occupied regions are related to cardiac cell differentiation. G. Top ten biological process GOs associated with TEAD1-GATA4 co-occupied regions (without CHD4) are involved with distinct biological processes, including vascular development.

To explore this possibility, we investigated the combined effect of TEAD and tissue-specific regulators at developmental enhancers. GATA motif is recognised by six family members, three of which, GATA4, GATA5, and GATA6, are highly expressed in cardiac tissues. To identify high-confident instances of GATA-TEAD co-binding in vivo, we overlapped human ventricle-specific H3K27ac bins with TEAD1 (35), GATA4 binding in mouse embryonic heart (32), and GATA6 binding in mouse posterior pharyngeal arches (37). Notably, de novo motif discovery performed on GATA6 peaks in the pharyngeal arches, (which contribute to the anterior pole of the heart) identifies TEAD amongst the top enriched motifs (37). We prioritised regions 1) shared across the three datasets; 2) containing TEAD and GATA motifs within a distance of 100nt and; 3) associated with genes with cardiac phenotypes (Fig 6A). We obtained five high-confident ventricle enhancers, four of which also intersected with CHD4 ChIP-seq peaks in the embryonic mouse ventricle. All the selected enhancers, cloned upstream of a minimal promoter and reporter gene, respond to their tissue-specific activator (GATA6 and GATA4) (Fig 6B and Fig S5AB). In all cases tested, addition of TEAD1 attenuates enhancer activation (Fig 6B and Fig S5A). Levels of inhibition by TEAD1 varied across enhancers (Fig S5C). We observed an additional decrease in enhancer activation when YAP and TEAD were co-transfected together (Fig 6B). In contrast, disruption of the TEAD-YAP complex (88), resulted in higher GATA activation for all the enhancers tested (Fig 6C), consistent with the presence of endogenous nuclear TEAD/YAP in 3T3 cells (Fig S5D). The identification of TEAD motifs close to tissue-specific TFs in developmental enhancers suggest that TEAD/YAP complex exerts its effect by binding to the enhancer. To address this, we generated mutant versions of one cardiac enhancer (Fig 6D). We observed that the repressive effect of TEAD/YAP complex is impaired in the absence of a canonical TEAD motif (Fig 6E), suggesting that TEAD/YAP ability to modulate enhancer activity is dependent on TEAD binding to the enhancer. Ruling out transcriptional repression as a generalised effect of our experimental system, TEAD and YAP increase luciferase activity driven by the *CCN2* promoter, a well-established TEAD target (Fig S5E). Finally, we assessed CHD4 occupancy at the *RAMP1* enhancer. Supporting a role for TEAD/YAP in recruiting CHD4 to this enhancer, we observed higher levels of CHD4 at the *RAMP1* enhancer when GATA6 was transfected together with TEAD/YAP complex, compared to GATA6 alone (Fig 6F).

**Figure 6.**
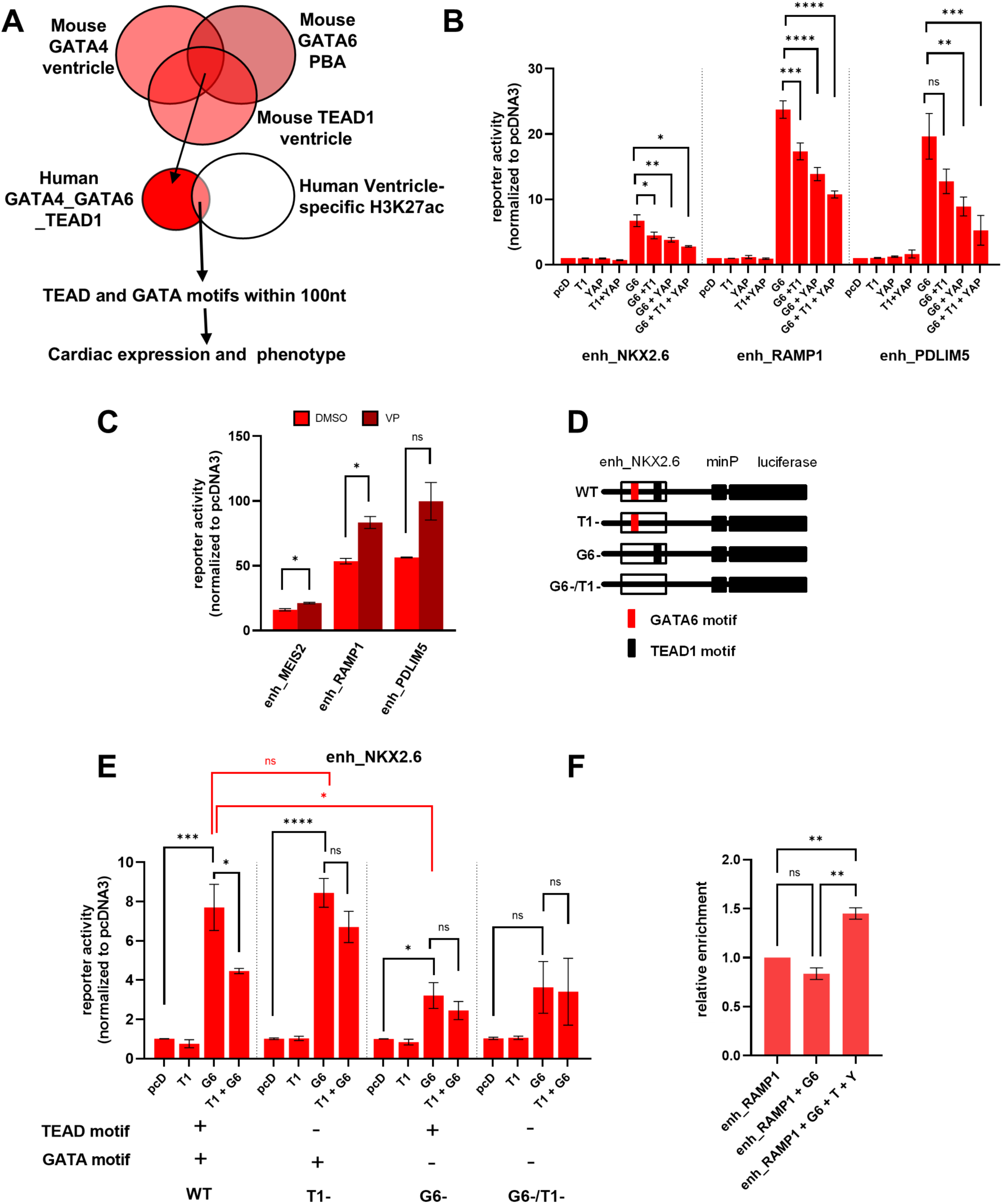
TEAD1 binding attenuates enhancer activation by cardiac-specific TFs. A. Workflow to identify high-confident candidate ventricle enhancers bound by TEAD1 and GATA4/GATA6. Mouse regions co-occupied by TEAD1, GATA4 and GATA6 were converted to human (hg38) coordinates. B. Luciferase activity driven by three ventricle-specific enhancers (enh_PDLIM5, enh_NKX2.6, enh_RAMP1) co-transfected with TEAD1 alone (T1), Gata6 alone (G6), YAP alone, Gata6 and TEAD1 (G6 + T1), TEAD1 and YAP (T1 + YAP), G6 and YAP, and G6, T1 and YAP in NIH3T3 cells, normalised against basal enhancer activity driven by an empty pcDNA3 vector (pcD). Co-expression of GATA6, YAP and TEAD1 significantly reduces enhancer activity. C. Luciferase activity driven by enh_MEIS2, enh_RAMP1 and enh_PDLIM5 co-transfected with GATA6 in NIH3T3 cells and treated with verteporfin (VP) or DMSO, normalised against basal enhancer level driven by an empty pcDNA3 vector (pcD). Inhibiting formation of the TEAD/YAP complex increases GATA6-dependent enhancer activation. Values are presented as the mean +/-SEM of two biological replicates, each obtained from the median of three experimental replicates. Asterisk denotes significant results. Significance was calculated using an unpaired t-test, * = P < 0.05. D. Diagram of the wild-type and mutant versions of enh_NKX2.6 in pGL4 plasmid (WT, G6-, T1– and G6-/T1-). E. Luciferase activity driven by wild-type enh_NKX2.6 and enh_NKX2.6 mutants. Each enh_NKX2.6 plasmids was co-transfected with TEAD1 alone (T1), with Gata6 alone (G6) and with Gata6 and TEAD1 (G6 + T1) in NIH3T3 cells. Luciferase values normalised against basal enhancer activity driven by an empty pcDNA3 vector (pcD). Compared to the wild-type enhancer, TEAD1 failed to significantly affect Gata6-induced enhancer activity upon mutation of TEAD1 binding site. Values (in B, E) are presented as the mean +/-SEM of three biological replicates, each obtained from the median of three experimental replicates. Asterisk denotes significant results. Significance was calculated using a one-way ANOVA with multiple comparisons for each of the enhancer constructs, **** = P < 0.0001, *** = P < 0.001, ** = P < 0.01, * = P < 0.05, ns = P > 0.05. H. Enrichment of CHD4 on_RAMP1 enhancer in the presence of GATA6 alone or GATA6, TEAD1 and YAP in NIH3T3 cells. All values are normalised against CHD4 enrichment on RAMP1 enhancer. Values are presented as the mean +/-SEM of two biological replicates, each obtained from the median of three experimental replicates. Significance was calculated using a one-way ANOVA with multiple comparisons for each of the enhancer constructs, **** = P < 0.0001, *** = P < 0.001, ** = P < 0.01, * = P < 0.05, ns = P > 0.05.

To generalise our findings, we chose a co-occurring partner of TEAD in a different tissue, CRX in the RPE (Fig 3C). For CRX-TEAD, given the lack of available CRX and TEAD1 binding information in the RPE, we selected high-confident RPE enhancers using RPE-specific H3K27ac bins and CRX peaks in the retina as a proxy, and association with genes expressed in the visual system. Three enhancers were prioritised for downstream analysis, using less stringent selection criteria than those adopted for ventricle enhancers (Fig 7A). We found that in all cases tested, addition of TEAD1 attenuates enhancer activation by CRX (Fig 7B). Again, levels of inhibition by TEAD1 varied across enhancers (Fig 7C). A mutant version of one RPE enhancer (Fig 7D), similarly to the mutated cardiac enhancer, showed that the repressive effect of TEAD/YAP is impaired in the absence of a TEAD canonical site (Fig 7E). Overall, these results suggest that TEAD/YAP binding at developmental enhancers can negatively interfere with their activation by tissue-specific TFs and uncover a broad, so far unrecognised, effect of TEAD on cell type-specific transcriptional programs (Fig 7F).

**Figure 7.**
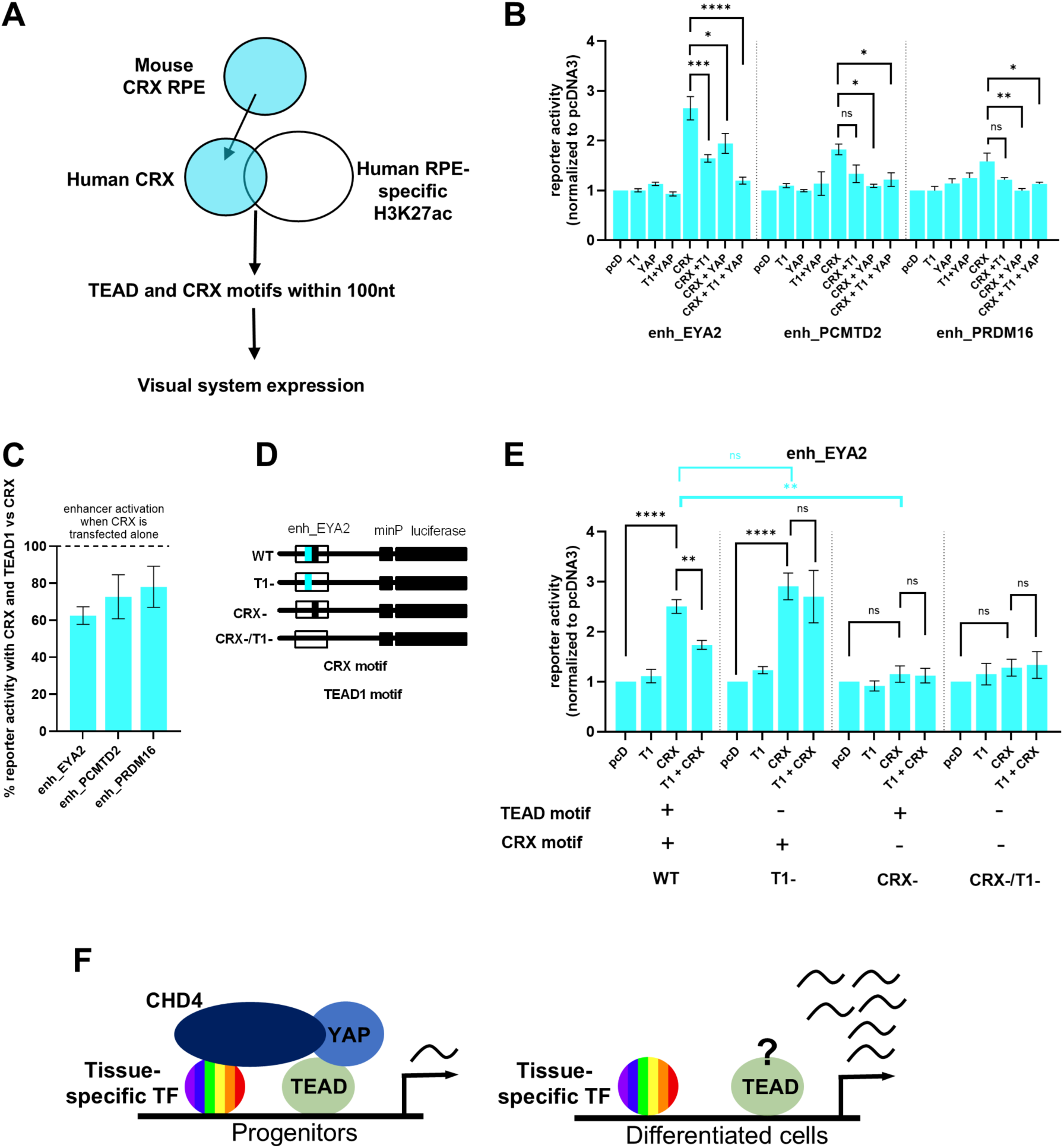
TEAD1 binding attenuates enhancer activation by RPE-specific TFs. A. Workflow followed to identify high-confident candidate RPE enhancers bound by TEAD1 and CRX. Mouse regions occupied by CRX were converted to human (hg38) coordinates. B. Luciferase activity driven by three RPE-specific enhancers (enh_EYA2, enh_PCMTD2 and enh_PRDM16) co-transfected with TEAD1 alone (T1), CRX alone, YAP alone, CRX and TEAD1 (CRX + T1), TEAD1 and YAP (T1 + YAP), CRX and YAP, and CRX, TEAD1 and YAP (CRX + T1 + YAP) in NIH3T3 cells, normalised against basal enhancer activity driven by an empty pcDNA3 vector (pcD). Co-expression of CRX, YAP and TEAD1 reduces enhancer activity to almost pcDNA3 levels. C. Percentage of CRX-induced luciferase reporter activity when CRX was overexpressed with TEAD1 compared to CRX overexpression alone. Percentages were calculated using the mean of three biological replicates. D. Diagram of the wild-type and mutant versions of enh_EYA2 pGL4 plasmid constructs (WT, CRX-, T1– and CRX-/T1-). E. Luciferase activity driven by wild-type enh_EYA2 and three enh_EYA2 mutants: T1-, CRX/T1– and CRX. Each of the four _EYA2 plasmids was co-transfected with TEAD1 alone (T1), with Crx alone (CRX) and with CRX and TEAD1 (CRX + T1) in NIH3T3 cells, normalised against basal enhancer activity driven by an empty pcDNA3 vector (pcD). When TEAD1 binding site is mutated, there is no significant inhibition of Crx-induced enhancer activity. Values (in B, E) are presented as the mean +/-SEM of 3 biological replicates, each obtained from the median of 3 experimental replicates. Asterisk denotes significant results. Significance was calculated using a one-way ANOVA with Tukey test for multiple comparisons of the means calculated for each enhancer plasmid (significance results are shown in black). In addition, unpaired t-tests were used to compare two conditions across enhancers (significance results shown in red/cyan), **** = P < 0.0001, ** = P < 0.01, * = P < 0.05, ns = P > 0.05. F. Working model. In progenitor cells, TEAD alongside YAP binds tissue-specific enhancers nearby tissue-specific activators and attenuates enhancer activity, reducing the expression of tissue-specific genes and slowing down cell differentiation. We propose that attenuation of enhancer activity is due to recruitment of CHD4 by the TEAD/YAP complex and tissue-specific activators. Reduced activity of TEAD in differentiated cells, possibly due to either TEAD no longer binding to the enhancer, or to its inability to recruit CHD4, results in full enhancer activation and sustained expression of tissue-specific genes. This mechanism could be part of a wider role of TEAD/YAP to slow down differentiation and maintain appropriate pools of progenitor-like cells to achieve correct organ size during organogenesis.

## DISCUSSION

Gene expression is largely controlled by TFs, which determine the global state of the cell. TF cooperativity is well established and widespread across TF networks. However, our knowledge of TF combinatorial binding largely derives from specific contexts (e.g. cell types or developmental time points). Here we provide a global overview of tissue-specific TF combinations at active enhancers in the developing human embryo. We present a range of evidence that supports our ability to detect functional co-occurring TF motifs. There are, however, limitations to using bioinformatics methods that rely on PWMs to predict functional binding events. The main one is that co-expressed TFs belonging to the same TF family recognise the same or very similar sites, limiting the accuracy of the predictions.

Our analysis identifies universal patterns of functional connectivity, shared by enhancers active in distinct tissues. A key finding is the recurrent co-occurrence of ubiquitous and tissue-specific TF motifs in active developmental enhancers. We find that tissue-specific enhancers from various, unrelated tissues are significantly enriched in motifs for the ubiquitous TFs TEAD, located near motifs recognised by tissue-specific TFs. TEAD TFs lack an activation domain and serve as a binding platform for coactivators, particularly YAP/TAZ. The hippo pathway regulates the nuclear-cytoplasmic shuttling of YAP and the formation of the TEAD-YAP complex (89). During mouse organogenesis, hippo signalling regulates cell proliferation and organ growth (90,91) across multiple organs, including the heart, craniofacial structures, lung, eye, brain, kidney and bile duct (92). Consistently, we identify enriched TEAD motifs in developmental enhancers active in the heart ventricle, palate, RPE, brain, kidney, lower limb and pancreas. By testing a subset of cardiac and RPE developmental enhancers, we find that TEAD1, together with its coactivator YAP, attenuates tissue-specific enhancer activation. The TEAD/YAP complex is widely recognized for its role as a transcriptional activator, yet it can also function as a transcriptional repressor through various mechanisms (34,93,94), including the recruitment of CHD4 and the NuRD complex (86). The preferential binding of TEAD1 with cardiac regulators at non-acetylated regions occupied by CHD4, suggests that combinatorial binding of TEAD1 with cardiac-enriched TFs attenuates enhancer activity via CHD4 recruitment. Supporting this hypothesis, TEAD1 repressive function in vitro relies on the presence of an activator; the TEAD/YAP complex does not repress enhancer activity per se and requires TEAD binding to DNA for this effect. Alongside the observed reduction in enhancer activity upon adding TEAD-YAP to transfected enhancers, CHD4 levels are increased at these enhancers. While our results point to cooperation of tissue-specific regulators with TEAD to recruit CHD4, the molecular contribution of each TFs to CHD4 recruitment, as well as the temporal dynamics of this process, are unclear. Additional studies on developmental enhancers in their genomic context, are required to fully elucidate TEAD role as a repressor.

The TEAD/YAP complex typically functions at early differentiation stages, in progenitor cells, and becomes downregulated at terminal stages of differentiation (95). In addition to regulating cell proliferation, this complex also impacts cell differentiation: loss of function experiments targeting TEAD or YAP result in decreased cell proliferation and rapid differentiation (93) while gain of function experiments produce the opposite effect – aberrant differentiation and increased proliferation (34,91,96–98). Our findings suggest a broad role for TEAD in regulating type-specific transcriptional programs. We propose that TEAD TFs, while promoting proliferation of precursor cells, also restrain their differentiation by recruiting CHD4, in collaboration with tissue-specific regulators, to lineage-specific enhancers (Fig 7F). Supporting this model, cardiac genomic regions co-occupied by GATA4, CHD4, and TEAD1 are primarily associated with the differentiation of cardiomyocytes, the main cell type in the heart. This suggests that TEAD1 may attenuate the activity of enhancers that control cardiomyocyte differentiation. In a developing organism, the timely differentiation of proliferating progenitors is essential for proper organ composition and size. Therefore, the attenuation of cell type differentiation programs by TEAD TFs may be a vital mechanism to prevent premature differentiation of progenitors ensuring the production of an adequate number of differentiated cells and contributing to appropriate organ size.

Understanding general principles of transcriptional networks is essential for deciphering how cell states are established and maintained, as well as for identifying the precise sequence codes that determine when and where genes are expressed. Beyond TEAD, we also identified the co-occurrence of other motif clusters recognized by ubiquitous TF families, such as TALE and ETS, alongside ‘First Search’, tissue-specific TFs. Our observations suggest that ubiquitously expressed TFs play a crucial role in shaping the chromatin landscape of developmental enhancers during human organogenesis.

The main focus of tissue-specific acquisition research revolves around the expression of tissue-specific TFs (99). Tissue-specific TF expression is central to stem cell differentiation protocols, reprogramming protocols and in developmental disease diagnostics. Our findings reveal that non-tissue-specific TFs may also contribute to tissue-specific gene regulation. Consistently, analysis of co-occurring motifs enriched in promoters of differentially expressed genes uncovered a potential involvement of non-tissue-specific TFs in driving tissue specificity (100). Given that enhancers have higher tissue specificity than promoters (101), the recurring enrichment of ubiquitous TF at tissue-specific enhancers highlights the significance of these TFs in tissue-specific transcriptional regulation. This insight raises important questions about the role of ubiquitous TFs in cell differentiation, lineage determination, and their overall impact on the acquisition of tissue specificity.

## DATA AVAILABILITY

Scripts and additional analysis scripts can be found in https://github.com/AGarciaMora/CocoTF.

## SUPPLEMENTARY DATA

SupplementaryFigures1-5.pdf

SupplementaryTables1-4.xlxs

## ACKNOWLEDGEMENTS

The authors would like to thank members of the Bobola lab for their scientific and technical insights, and the Bioimaging Facility at the University of Manchester for help with the imaging performed in this project.

## FUNDING

This work was supported by a Medical Research Council (MRC) (http://www.mrc.ukri.org) studentship to AGM, and Biotechnology and Biological Sciences Research Council (BBSRC) (http://www.bbsrc.ukri.org) grants BB/N00907X/1 and BB/T007761/1 to NB. The Bioimaging Facility microscopes used in this study were purchased with grants from BBSRC, Wellcome and the University of Manchester Strategic Fund.

## CONFLICT OF INTEREST

No conflict of interest exists that might raise the question of bias in the work reported or the conclusions, implications, or opinions stated.

## Notes

### Competing Interest Statement

The authors have declared no competing interest.

### Summary of Updates

The result section is expanded in the more recent version; this has also resulted in major modifications of figures 2-7. We have also added supplemental figures.

