## Supplemental Figures for "Systematic analysis of transcription factor combinatorial binding uncovers TEAD1 as an antagonist of tissue-specific transcription factors in human organogenesis"

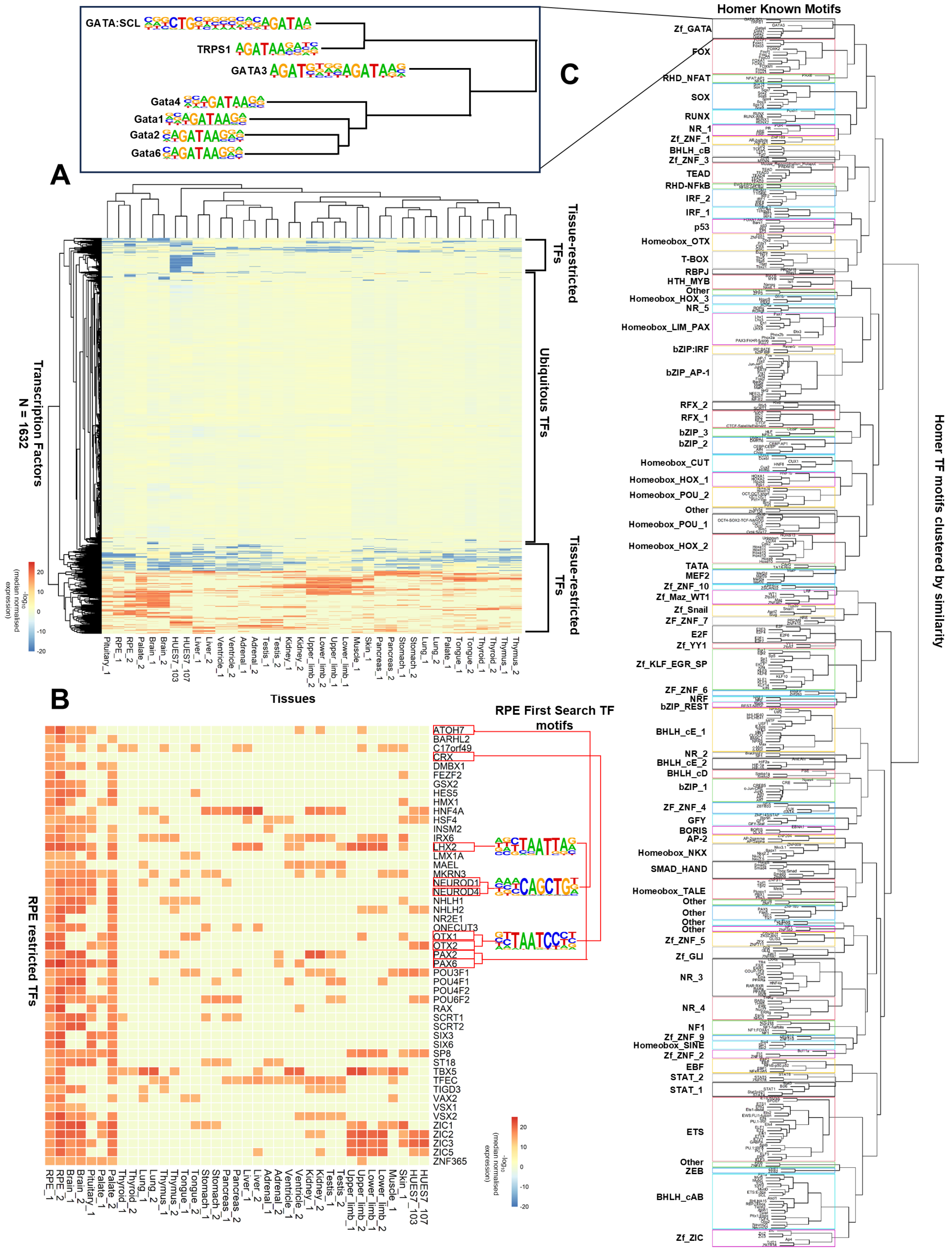


**Figure S1 (Supplementary to Figure 1).** A. List of TFs from (5) represented in the RNA-seq data from (30). All tissues were duplicated except for pituitary, muscle and skin. Raw counts were down sampled based on the 75th percentile, expression of each TF was divided by their median expression across all tissues and log median-normalised expression was used to plot the heatmap. Hierarchical clustering was performed (k = 10). Ubiquitous TFs, with expression constant across tissues clustered in a large cluster labelled ubiquitous TF cluster. The remaining clusters were considered tissue-restricted. B. Heatmap showing RPE-restricted TFs. TFs with expression higher than 150 times their median expression in one tissue were considered restricted for that tissue. Those RPE-restricted TFs, whose recognition motifs (or a motif belonging to a member of the same cluster seen in Fig S1C) are within the top 20 motif enrichment analysis of RPE-specific H3K27ac 1kb bins were considered 'First Search TFs'. Nine tissue-restricted TFs, ATOH7, CRX, LHX2, PAX2, PAX6, NEUROD1, NEUROD4, OTX1 AND OTX2, had a motif represented in the top 20 RPE motif enrichment analysis results. However, due to the similarity of their motifs, these were collapsed to three 'First Search' TF motifs. C. Hierarchical clustering tree of all motifs contained in the HOMER motif library, grouped by motif similarity. Cut-off used was height = 0.5, which resulted in 79 clusters (details on motifs belonging to each cluster can also be found in Table S3). Each cluster was assigned a name based on its cognate TF families and subfamilies. If multiple TF motif families clustered together, a combination of family names was used. When classification based on subfamilies was not possible, numbers were assigned to the clusters.

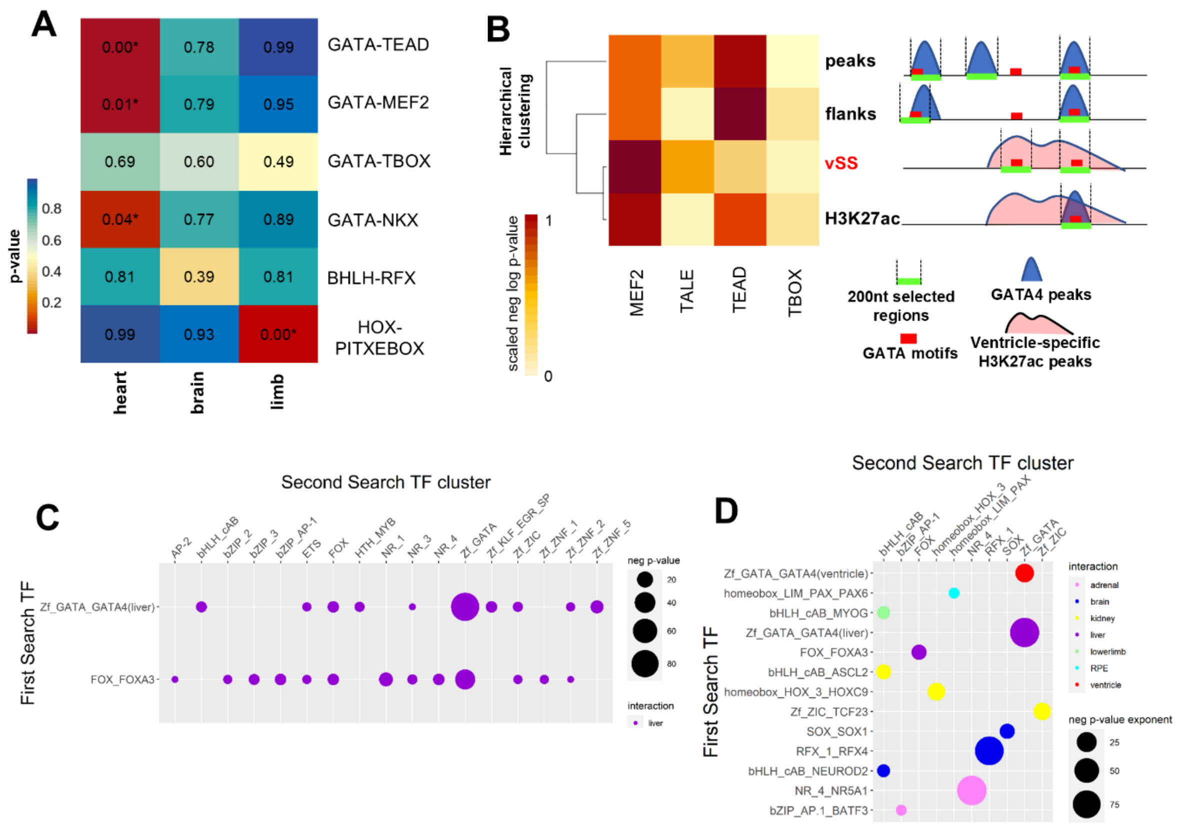


**Figure S2 (Supplementary to Figure 2).** A. Heatmap showing p-values of the probability of GATA-TEAD, GATA-MEF2, GATA-TBX, GATA-NKX, bHLH-RFX, HOX-PITXEBOX motif pairs co-occurring (within 100nt) in heart enhancers compared to brain and limb specific enhancers (from VISTA). P-values were calculated using a hypergeometric test and significant results ($p < 0.05$) are indicated with an asterisk. GATA-TBX pair was not enriched in the heart compared to brain and limb enhancers; however very few instances of TBX motifs were detected in the VISTA enhancers, which has an effect on hypergeometric p-values. B. Motif enrichment analysis of GATA4 mouse heart ventricle ChIP-seq validates the pipeline's ventricle results. Three datasets were analysed and compared to ventricle 'Second Search' (vSS) results using GATA in the ventricle as a 'First Search' TF. Dataset 'peaks' consists of 200nt GATA4 ChIP-seq peaks, dataset "flanks" consists of 100nt flanks either side of any GATA motifs found within GATA4 'peaks' and dataset 'H3K27ac' consists of GATA4 'flanks' overlapping human ventricle-specific H3K27ac 1kb bins converted to the mouse genome. 'Second search TF' MEF2, TALE, TEAD and TBOX motifs are among the top 10 enriched motifs in all datasets. Heatmap shows enrichment p-values of MEF2, TALE, TEAD and TBOX motifs obtained from motif enrichment analysis of 'peaks, 'flanks', 'vSS' and 'H3K27ac' performed by HOMER. Neg log p-values are scaled across experiments, to show patterns of significance rather than absolute values. GATA motifs were excluded from the results as the GATA motifs was not masked in some of the motif enrichment analysis. C. Bubbleplot of liver results, focusing on the potential direct or indirect interactors of FOX and GATA TFs at liver-specific enhancers. FOX and GATA co-occur with distinct clusters of 'Second Search' motifs. D. Bubbleplot of homotypic motif co-occurrences, where 'First Search' and 'Second Search' TFs belong to the same cluster. Colours correspond to the ones assigned in Fig 1B. The size of the bubbles matches the negative log p-value: larger bubbles represent smaller p-values (in C,D).

**A**

**Heart-specific (relative to shared peaks)**

| ***De novo* motif** | **Enrichment** | **P-value** | **Match** |
|---|---|---|---|
| 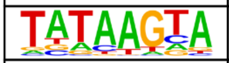 | 12% | 1e-1482 | **MEF2** |
| 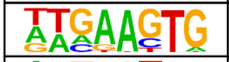 | 8% | 1e-1301 | **NKX2** |
| 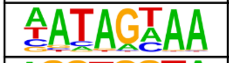 | 10% | 1e-1259 | **RFX** |
| 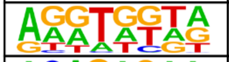 | 6% | 1e-1193 | **T-BOX** |
| 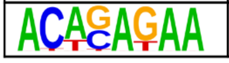 | 5% | 1e-1116 | **STAT** |

**B**

**Liver-specific (relative to shared peaks)**

| ***De novo* motif** | **Enrichment** | **P-value** | **Match** |
|---|---|---|---|
| 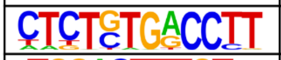 | 8% | 1e-223 | **NR** |
| 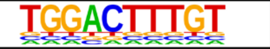 | 7% | 1e-196 | **HNF4** |
| 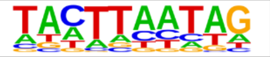 | 7% | 1e-191 | **Homeobox** |
| 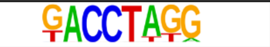 | 13% | 1e-181 | **NR** |
| 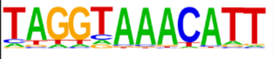 | 7% | 1e-181 | **Forkhead** |

**C**

**keratinocyte-specific (relative to shared peaks)**

| ***De novo* motif** | **Enrichment** | **P-value** | **Match** |
|---|---|---|---|
| 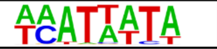 | 11% | 1e-1661 | **SOX** |
| 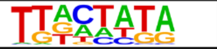 | 9% | 1e-1521 | **POU** |
| 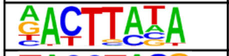 | 10% | 1e-1366 | **NKX3** |
| 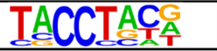 | 8% | 1e-1338 | **ZEB** |
| 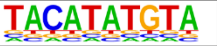 | 9% | 1e-1278 | **TWIST** |

**D**

**enriched in shared peaks (relative to tissue-specific)**

| ***De novo* motif** | **Enrichment** | **P-value** | **Match** |
|---|---|---|---|
| 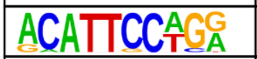 | 82% | 1e-48 | **TEAD** |
| 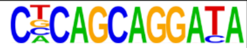 | 1% | 1e-12 | **ZIC** |

**Figure S3 (Supplementary to Figure 4).** A. Top five enriched motifs in TEAD1 ventricle-specific peaks relative to shared peaks. Within the top five results are motifs matching heart-specific TFs such as NKX, T-BOX and MEF2. B. Top five enriched motifs in TEAD1 liver-specific peaks relative to shared peaks. Within the top five results are motifs matching liver-specific TFs, such as HNF4, and nuclear receptor. C. Top five enriched motifs in TEAD1 keratinocyte-specific peaks relative to shared peaks. Within the top five motifs are important TFs for skin differentiation belonging to the POU family. D. Enriched HOMER de novo motifs in TEAD1 non-tissue-specific peaks. Liver-, ventricle- and keratinocyte-specific TEAD1 peaks were used as background. All p-values were obtained as part of the HOMER de novo output.

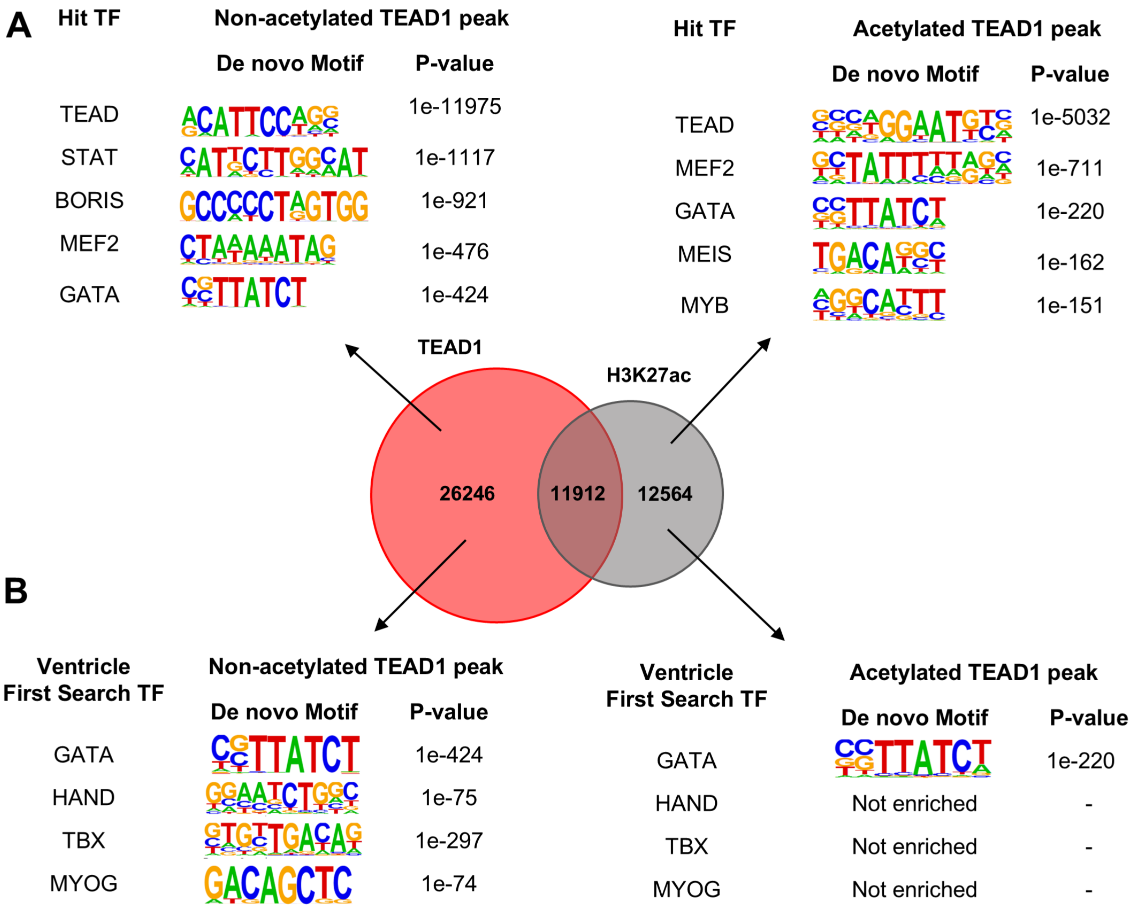


**Figure S4 (Supplementary to Figure 5).** A. Overlap between TEAD1 and H3K27ac ChIP-seq peaks in the embryonic heart ventricle and top 5 de novo motifs from HOMER motif enrichment analysis. B. Representation of cardiac 'First Search' TFs in HOMER de novo motif enrichment analysis of TEAD1 peaks overlapping with H3K27ac and without H3K27ac. Motifs bound by tissue restricted TFs are more enriched in non-H3K27ac TEAD peaks. Except for GATA, no 'First Search' TF motif is enriched in TEAD1 acetylated peaks.

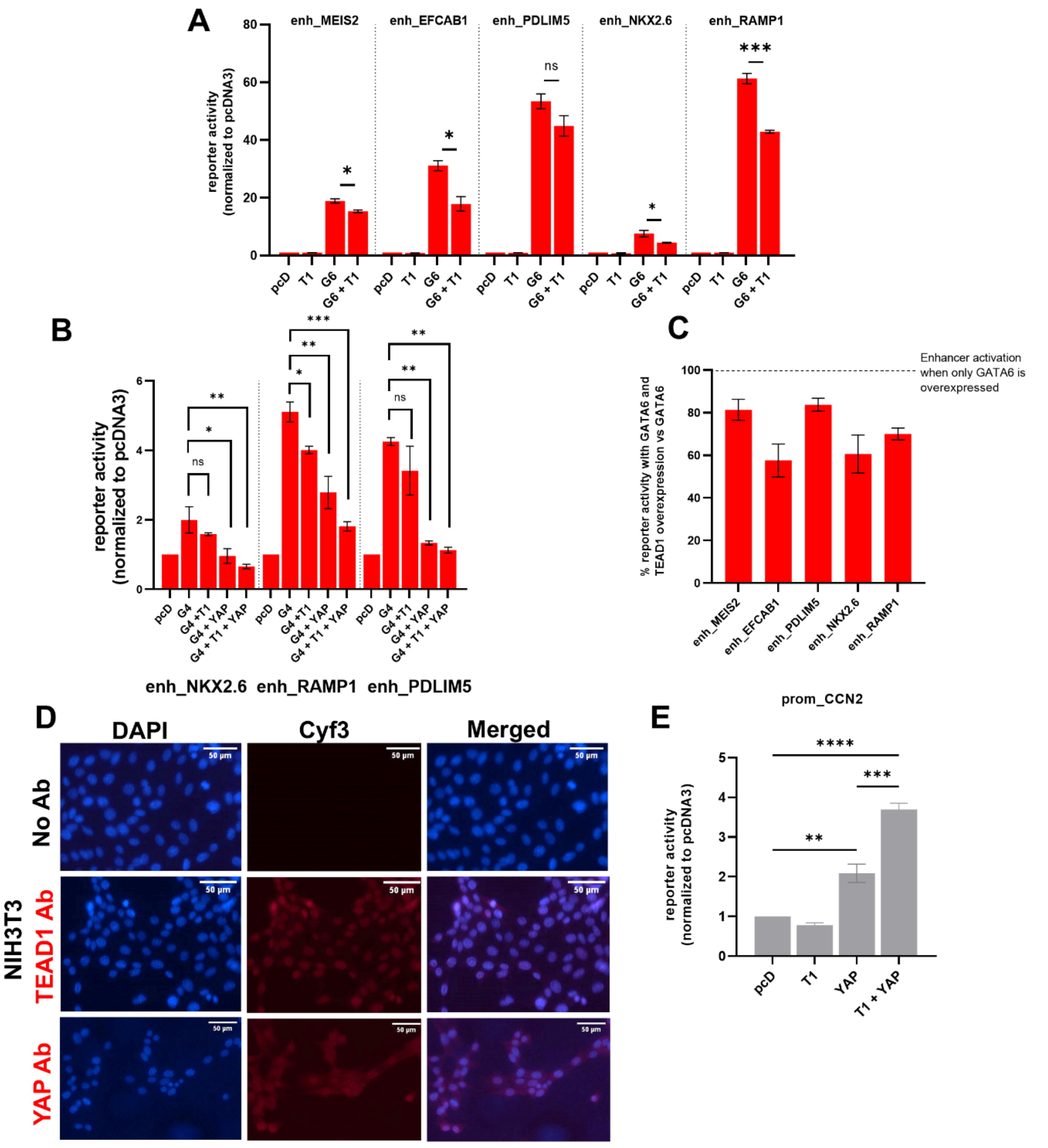


**Figure S5 (Supplementary to Figure 5).** A. Luciferase reporter activity driven by five ventricle-specific enhancers (enh_MEIS2, enh_EFCAB1, enh_PDLIM5, enh_NKX2.6, enh_RAMP1) co-transfected with TEAD1 alone (T1), with GATA6 alone (G6) and with GATA6 and TEAD1 (G6 + T1) in NIH3T3 cells, normalised against basal enhancer levels driven by an empty pcDNA3 vector (pcD). Gata6 significantly increases luciferase reporter activity in all five enhancers, and co-transfecting Gata6 and TEAD1 significantly reduces Gata6-induced enhancer activity in all enhancers except for enh_PDLIM5. Values are presented as the mean +/- SEM of three biological replicates, each representing the median of three experimental replicates. Asterisk denotes significant results. Significance was calculated using an unpaired t-test comparing luciferase activation under G6 and G6 + T1 condition in each of the enhancer constructs, **** = $P < 0.0001$, ** = $P < 0.01$, * = $P < 0.05$, ns = $P > 0.05$. B. Luciferase reporter activity driven by three ventricle-specific enhancers (enh_NKX2.6, enh_RAMP1, enh_PDLIM5) co-transfected with GATA4 alone (G4), with GATA4 and TEAD1 (G4 + T1), with GATA4 and YAP (G4 + YAP) and with

GATA4, TEAD1 and YAP (G4 + T1 + YAP) in NIH3T3 cells, normalised against basal enhancer levels driven by an empty pcDNA3 vector (pcD). GATA4 increases luciferase reporter activity in all three enhancers, and co-transfecting GATA4 with TEAD1 and YAP significantly reduces GATA4-induced enhancer activity in all enhancers tested enhancers. Values are presented as the mean +/- SEM of two biological replicates, each representing the median of three experimental replicates. C. Percentage of GATA6-induced luciferase reporter activity when Gata6 was overexpressed with TEAD1 compared to Gata6 overexpression alone. Percentages were calculated using the mean of three biological replicates. D. NIH3T3 cells incubated with YAP or TEAD1 antibody and a secondary antibody containing an Alexa 555 fluorophore. Whilst TEAD1 is concentrated in the nucleus, YAP can be found both in the cytoplasm and nucleus. E. Luciferase activity driven by the *Ccn2* promoter (prom_Ccn2) co-transfected with TEAD1 (T1) expression vector only, YAP expression vector only and TEAD1 and YAP (T1 + YAP), in NIH3T3 cells. Absolute values are normalised to luciferase activity measured with an empty vector (pcD), hence plotted values represent fold change. Significance (in B, E) was calculated using a one-way ANOVA with Tukey multiple comparisons of the means for each one of the promoter plasmids, **** = $P < 0.0001$, *** = $P < 0.001$, ** = $P < 0.01$, ns = $P > 0.05$.

**Table S1 (supplemental to Materials and Methods).** Primer sequences; mutated NKX2.6 and EYA2 enhancer sequences; conditions for luciferase assays.

**Table S2 (supplemental to Fig 1).** Complete list of tissue-restricted TFs. Tissue-restricted TFs, whose recognition motifs (or a motif belonging to a member of the same cluster seen in Fig S1C) are within the top 20 motif enrichment analysis for their corresponding tissue are shown in bold.

**Table S3 (supplemental to Fig 1).** Table obtained from hierarchical clustering tree of all motifs contained in the HOMER motif library, grouped by motif similarity. Each row shows the motifs encompassing each cluster, named according to the homer motif library nomenclature.

**Table S4 (supplemental to Fig 1).** ‘First Search’ TFs for each tissue, with tissue-specific expression and with motifs enriched within the top 20 motif enrichment analysis results of their corresponding tissue-specific H3K27ac putative enhancers. TFs that bind the same TFBS are shown in the same row. The name of the chosen PWM as provided by HOMER is shown in column "top enriched motif". The highest expressed TF from each row is shown in bold. The cluster to which the motif belongs to is shown in column “motif cluster”. The assigned name for each ‘First Search’ TF in figure 2 consists of motif cluster + highest expressed TF.
